# Optimized two-color single-molecule tracking of fast-diffusing membrane receptors

**DOI:** 10.1101/2023.03.17.533099

**Authors:** Chiara Schirripa Spagnolo, Aldo Moscardini, Rosy Amodeo, Fabio Beltram, Stefano Luin

## Abstract

Single particle tracking (SPT) combined with total internal reflection fluorescence (TIRF) microscopy is an outstanding approach to decipher crucial molecular mechanisms on the cell membrane at the nanometric scale. In multicolor configurations it can even be the ideal tool to investigate interactions, but this is hindered by a number of experimental challenges. We systematically and quantitatively analyze the impact of all the necessary sub-elements of any SPT-TIRF setup on signal-to-noise ratio, especially in dynamic studies using minimally-invasive dyes for biomolecule labeling. We show that the dominant limiting factor is the autofluorescence originating from the commonly-used optical glass. We identify and test a different material and show a significant improvement of signal-to-noise ratio in a multichannel TIRF configuration employing the new glass covers. We also address the problem of photobleaching of fluorescent probes by presenting effective approaches suited to a multicolor implementation that requires simultaneous stabilization of multiple dyes. We apply the developed protocol to the analysis of p75 receptors labeled by two fluorophores on the membrane of living cells. Our strategy yields reliable, simultaneous two-color SPT, even for fast-diffusing receptors, enabling this study under conditions not accessible with standard experimental configurations. We argue that the present protocol can pave the way for multicolor super-resolved localization and tracking of single molecules by TIRF microscopy, much expanding the potential of SPT.

## 1. Introduction

Total internal reflection fluorescence (TIRF) microscopy is a powerful technique for visualizing key biological events occurring on the cell membrane with outstanding signal-to-noise ratio and spatial resolution [1–3]. TIRF microscopy can even reach the single-molecule level. Especially combined with single-particle tracking (SPT) this can yield an accurate description of the spatio-temporal organization of the cell membrane environment and provide much information on its structure and functioning [4–6]. SPT provides the exceptional ability to observe directly each molecule at work in a living system. So far, the major developments reported are in the context of single-color SPT, where a single particle type is labelled and tracked over time. These studies provided many new insights into molecule dynamics and homo-interactions, revealing the strong correlation between these and biological mechanisms [7–9].

Multicolor extensions of TIRF-based SPT allow the quantitative study, at the single-molecule level, of hetero-interactions on the cell membrane, e.g. interaction of different receptors with one another or of receptors with their ligands. Indeed, signal transduction is typically determined by complex interplays of different molecules and deciphering such mechanisms requires the simultaneous observation of different moieties.

However, multicolor SPT applications are still limited and are hindered by several experimental difficulties.

From the point of view of the microscopy setup, the simplest implementations are based on the use of Quantum Dots (Qdots), which, therefore, are typically the fluorescent labels of choice [10–12]. Indeed, Qdots are convenient for SPT, in comparison to other labels such as organic dyes, owing to their high brightness and photostability, which ensure optimal tracking performance (i.e. high localization accuracy, extended tracking). Additional advantages of Qdots for multicolor studies include narrow emission spectra, allowing for good channel separation with minimal crosstalk, and the possibility to use a single excitation, resulting in a simpler setup but also helping in minimizing induced autofluorescence and background in general [13,14]. A drawback of Qdots is their relatively large size, with a typical reported diameter of 10-50 nm, compared to 1-2 nm for organic dyes [13,15]. Indeed, some studies revealed the impact of steric hindrance and a slowed dynamics due to Qdot labelling. Marchetti et al. observed a radically slower profile of the diffusion coefficient of p75^NTR^ receptors labelled with Qdots compared to the same receptors labelled with organic dyes [7]. Similarly, Abraham et al. observed that, in comparison to the case of labelling by small organic dyes, Qdots hinder B cell receptors mobility [16]. The observed large size of Qdots stems not only from their intrinsic size, but also from the need of additional molecules (antibodies or streptavidin) required to functionalize them for labeling applications. Alternative strategies for Qdot labeling without such large molecules still require improvements; in any case, to date, it is difficult to achieve control of both stoichiometry and efficiency in labeling with Qdots, in contrast to organic dyes for which several protocols are available for minimally-invasive, stoichiometry-controlled and efficient labelling [14].

Here we investigate the experimental challenges related to two-color single-particle tracking using small organic dyes, a technique that we implemented using a TIRF microscope by adding a laser combiner before the optical fiber coupled to the setup and DualView to acquire two emission channels simultaneously on a single fast electron-multiplier CCD. We analyze the two critical issues related to the choice of these labels: low signal-to-noise ratio and photobleaching. The former is caused by the relatively low brightness of the dyes and the background that can be significant especially with the multiple excitations required for these dyes. We show that several background sources exist amongst the necessary elements of a TIRF system, and their unwanted emission becomes more severe in multicolor implementations. Limited reports exist on these effects. For example, there are some studies of autofluorescence from optical glasses or cells but they are few and mainly concern excitation in the UV range [17–21]. In particular, here we observe a strong impact of the background arising from standard cover glasses, and we describe the use of a material that significantly improves the final signal-to-noise ratio for the case of dyes in a two-color configuration.

We also deal with photobleaching. This is a complex phenomenon caused by the combination of different mechanisms; several strategies for reducing its impact were proposed, but the best solution strongly depends on the considered dye [22,23]. Therefore, reducing photobleaching in multicolor studies is even more challenging because a common solution must be found for different dyes. We shall present an effective strategy to reduce photobleaching for the two dyes we select.

Finally, we shall show that the approach presented here makes it possible to perform two-color SPT experiments in living cells on fast p75^NTR^ receptors labelled with two different organic dyes, obtaining reliable tracking results in both channels.

## 2. Results

### 2.1 Effects of background on the SNR of single molecules

We measured the background in a TIRF system without any fluorescent sample, but with all the other essential elements (immersion oil, glass coverslip bottom of the dishes, water medium; Figure 1A). This background depends strongly on excitation-and detection-band combinations, as observed in typical configurations used for single-molecule fluorescence (Figure 1A). These phenomena strongly impact on the SNR of dyes, especially in single-molecule measurements. We measured the SNR of the dyes Atto 565 and Abberior STAR 635p: they have identical brightness, comparable extinction coefficient, and quantum yield (Table S1), but the SNR of the second resulted significantly smaller, because of the different background associated with the individual excitation and detection wavelengths (Figure 1B). We observed this behavior both on immobile dyes adherent to the thin glass bottom of Petri dishes for microscopy (called “cover glass” from here on for simplicity) and on dyes labelling the neurotrophic TrkA receptors that diffused on the membrane of living cells. As expected, for each dye, the SNR was higher when the dyes adhered to the coverslip than when they labelled TrkA receptors. This can be explained by an additional biological source of autofluorescence, by the slightly lower signal caused by the distance of the membrane from the glass-water interface, and by the impact of the receptor movement: diffusion causes a degradation of the SNR, as we also observed by comparing the SNR of two receptors having different diffusivities, TrkA and P75^NTR^ (Figure S1).

**Figure 1.**
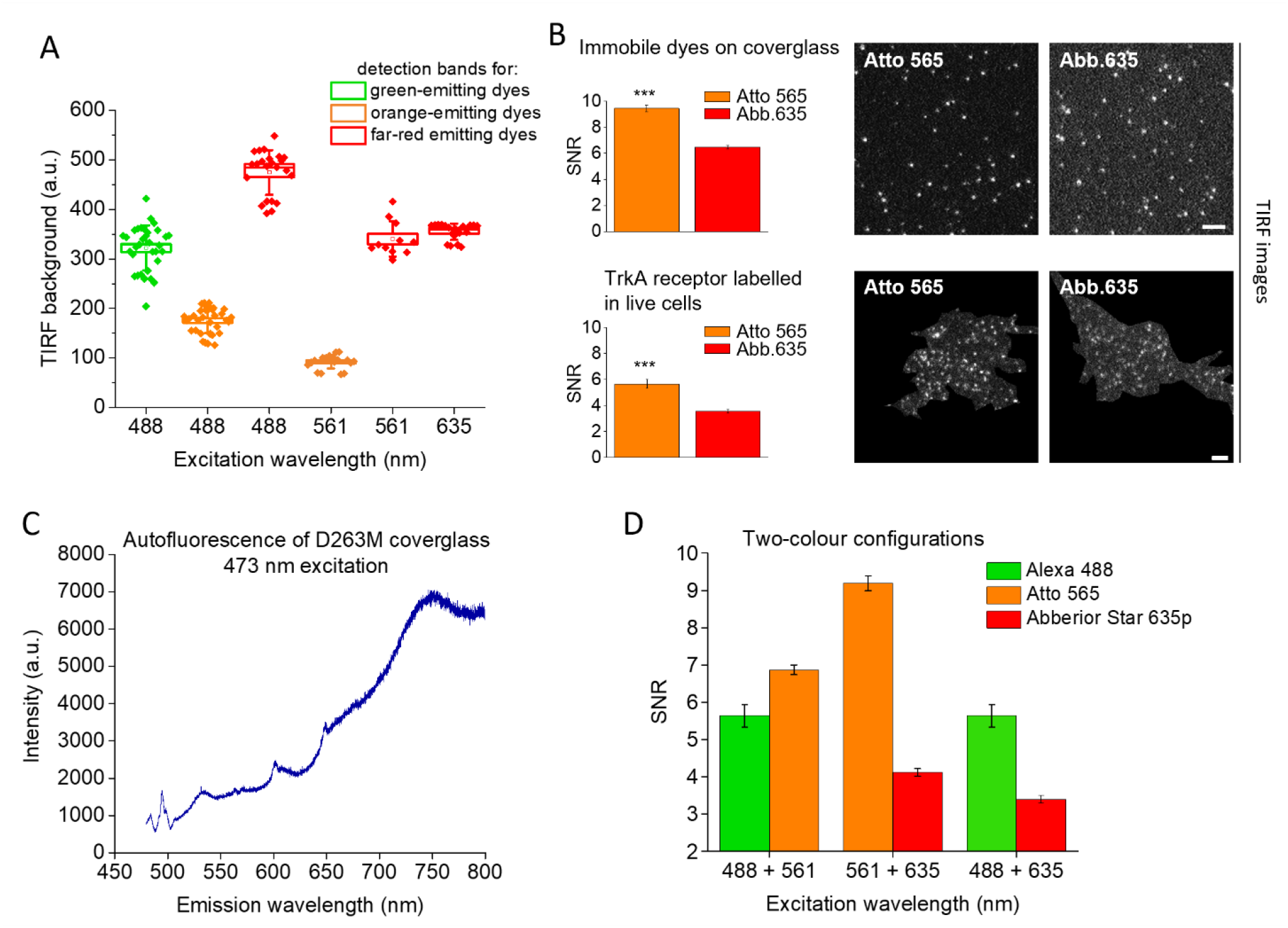
Background behavior and its effects in single-molecule TIRF microscopy. A) Background detected in TIRF with different combinations of excitation and detection bands with standard glass (D263M)-bottom Petri dishes (at fixed excitation power and CCD parameters). Bands and parameters are detailed in Materials and Methods and Supplementary Methods sections. Box: SEM; whisker: SD; dots: individual fields of view, from 5 independent repetitions. B) Signal-to-noise ratio (SNR) measured in single-molecule TIRF for the two dyes Atto 565 and Abberior STAR 635p (Abb.635) having comparable brightness (see Supplementary Table 1) but excitation and detection bands corresponding to different background levels. Results reported for dyes adherent to cover glass (top) and labelling moving TrkA receptors on the membrane of adherent live SH-SY5Y cells (bottom), with corresponding representative TIRF images on the right (the region outside the basal cell membrane is blackened for images on the bottom). Data are mean ± SEM. Dyes on cover glass: 7 movies in different fields of view from 2 independent repetitions; dyes labelling TrkA receptors: 16 movies in different fields of view from 3 independent repetitions. Intensity signal mediated in space and time on a total of 2000-20000 spots for each movie. *** P < 0.001, Student’s t-test. Scale bar: 3 mM C) Typical emission spectrum for D263M cover glass using excitation at 473 nm. D) Evaluation of SNR in three two-channel configurations (estimated from single-channel measurements as described in the Materials and Methods section), using dyes adherent to cover glass. Data are mean ± SEM, obtained from 4-9 movies in different fields of view from 2 independent repetitions; intensity signals mediated on 2000-13000 spots for each movie.

Figure 1A suggests that these effects may become even more relevant in a multicolor setup because of the high background in the far-red region upon excitation with a shorter wavelength such as 488 or 565 nm. We observed that a relevant contribution to this background is due to the emission of optical glass, in particular of the cover glass supporting the sample. Indeed, we measured the emission of a typical cover glass (Schott D263M) with a 473 nm excitation in a microRaman system and we observed higher emission in the longer-wavelength portion of the spectrum (Figure 1C).

We then compared three possible implementations of a two-color TIRF system with the following three dye pairs (Figure 1D, see also Materials and Methods section): (i) 488-excitation, green dye paired with a 561-excitation, orange dye; (ii) 561-excitation, orange dye paired with a 635-excitation red dye; (iii) 488-excitation, green dye paired with a 635-excitation red dye. We selected the dyes (Alexa 488, Atto 565 and Abberior STAR 635p) among the ones already used in single-molecule applications with the highest brightness for the corresponding channel [7,14,24].

We observed that the 488-565 couple was the one with the most homogeneous properties in the two channels; the 488-635 couple was the less advantageous; the 565-635 couple had a very different SNR between the two channels, with the first channel having more-than-double SNR. The choice of the best configuration may depend on the application of interest, e.g. on the diffusivity of the molecules imaged in the two channels. However, the 565-635 and 488-635 couples showed a considerable limit in the far-red channel owing to the background detected in that region. From these preliminary observations we concluded that (i) the behavior of the background caused by the necessary elements in a TIRF system strongly influences the SNR of single molecules; (ii) in the choice of channels for single and multi-color studies it is important to consider the different background contributions associated with the various excitation-detection pairs rather than consider only the dye brightness; (iii) glass emission can be a limiting factor for multi-color applications.

### 2.2 N-PK51 optical glass as cover glass material for TIRF microscopy

Due to the high impact of the cover glass fluorescence on the background, especially for multicolor applications, we compared the emission intensity of different cover glasses at different excitations and emission bands combinations (Figure 2A).

**Figure 2.**
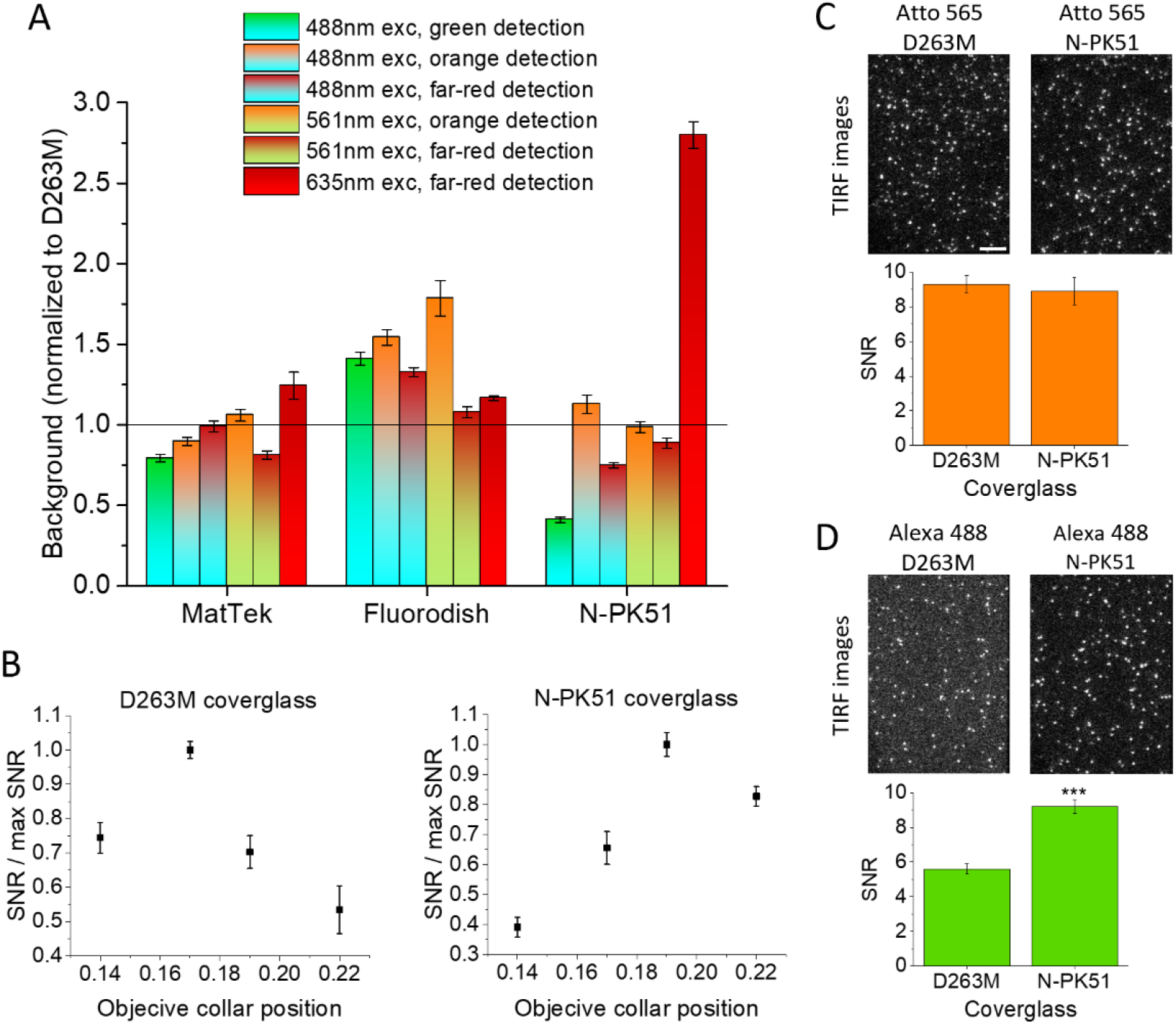
Cover glasses comparison and use of NPK51 glass for single-molecule TIRF imaging. A) Ratio between the background detected using different cover glass kinds and using D263M cover glass. Bar color gradients illustrate excitation (bottom) and detection (top) wavelengths. Data are mean ± SEM, obtained from 8-23 fields of view from 2 independent repetitions. B) SNR of single molecules of Atto 565 adhered on D263M (left) and NPK51 (right) cover glasses at different objective collar positions, normalized to the maximum observed SNR. Measurements were performed at 37 °C. Data are mean ± SEM, obtained from 5 movies in different fields of view from 2 independent repetitions, with 2000-20000 spots analyzed in each movie. C) Top: representative TIRF images of single molecules of Atto 565 adhered on D263M (left) and NPK51 (right) cover glass. Bottom: corresponding SNR. D) Top: representative TIRF images of single molecules of Alexa 488 adhered on D263M (left) and NPK51 (right) cover glass. Bottom: corresponding SNR. Images in C) and D) were acquired separately in single-channel configuration. Data are mean ± SEM, obtained from 10 movies in different fields of view from 2 independent repetitions, with 2000-20000 spots analyzed in each movie. *** P < 0.001, Student’s t-test. Scale bar: 5 mM; images reported with the same grayscale.

Schott D263M glass is used to make cover glasses of different brands and is incorporated in glass-bottom dishes from many companies (such as Willco Dishes, Ibidi, Azer Scientific). We identified a few brands that use different glasses for the bottom of their dishes, e.g. MatTek dish and FluoroDish. We also investigated the possibility of using different non-standard materials. TIRF systems, especially the objective-based commercial ones, place severe limitations on the optical properties of the cover glass. The setups are designed to be used with a cover glass that has a thickness of 0.17 mm and a refractive index of about 1.52 (around 550 nm). We considered Schott N-PK51 glass as a possible candidate (because of its refraction index and considering the few published data on glasses fluorescence; see Supplementary note and Figure S2) and obtained some custom cover glasses made from this material (specifications in Table S2). In the autofluorescence comparisons between MatTek, FluoroDish and N-PK51, we found the biggest improvement over the D263M case using N-PK51, especially considering the green-region emission under excitation at 488 nm, which was reduced to 40% compared to D263M. Instead, on N-PK51, the emission detected in the far-red under red excitation was 2.4 times higher compared to D263M, more than compensating the small improvement in the far-red emission upon blue excitation in a hypothetical 488-633 couple for simultaneous two-color applications. We did not find a material significantly improving the low SNRs for Abberior Star 635p in two-color configurations shown in Figure 1D; therefore, in the following we will mostly show results on dyes with excitations at 488 nm and 561 nm.

Modern objectives are equipped with a correction collar, which can compensate for small variations (a few micrometers) in cover glass thickness or variations in refractive indexes typically caused by temperature changes. We searched for the best position of the objective correction collar using the standard D263M cover glasses and the N-PK51 custom cover glasses. We measured the SNR of Atto 565 as a function of collar positions, with optimized focus at 37° C, as needed for live cell imaging (Figure 2B). As expected, the optimal position for D263M was that for 0.17 thickness. We found a small shift for NPK51, likely due to the small difference in refractive index (Table S2). The SNR for Atto 565 dye in the optimal positions was comparable between the two materials (Figure 2C); in fact, for its excitation and detection, we measured a similar autofluorescence of the two materials. In conclusion, we could correct the impact of the refractive index difference by using the correction collar.

When comparing the SNR for 488 dyes, we observed a significant improvement using the N-PK51 cover glass (Figure 2D). Thus, a first advantage of this material relates to single-color applications where 488 dyes are preferred, and the final obtainable SNR of these dyes, typically having lower brightness, is comparable to that achievable with brighter dyes emitting at different wavelengths (Table S1). Thanks to the significant improvement in the 488-dye channel, the material promises a better two-color performance compared to standard D263M using 488+565 channels.

### 2.3 Two-color single-molecule imaging: immersion oil

To investigate the performance of the N-PK51 glass in a simultaneous two-color setup, we employed a single-fiber excitation setup. We minimized chromatic aberration, in particular to obtain focal positions as close as possible in the two channels, starting with the observation that these kinds of aberrations are influenced by immersion oil (Figure 3 A,B,C). We searched for the best immersion oil in our system for N-PK51 and D263M and found different behaviors for the two materials (Figure 3 A,B,C) that probably stem from a difference in the Abbe number (Table S2). All the immersion oils have the same refractive index (Table S3); indeed, we found the same optimal collar position and negligible differences in maximum SNRs comparing different oils (Figure S3). They have some differences in their Abbe number (Table S3) but, interestingly, we found different degrees of chromatic aberration even among oils with the same theoretical Abbe number (e.g. Nikon and Olympus oils with the D263M cover glass, Figure 3B).

**Figure 3.**
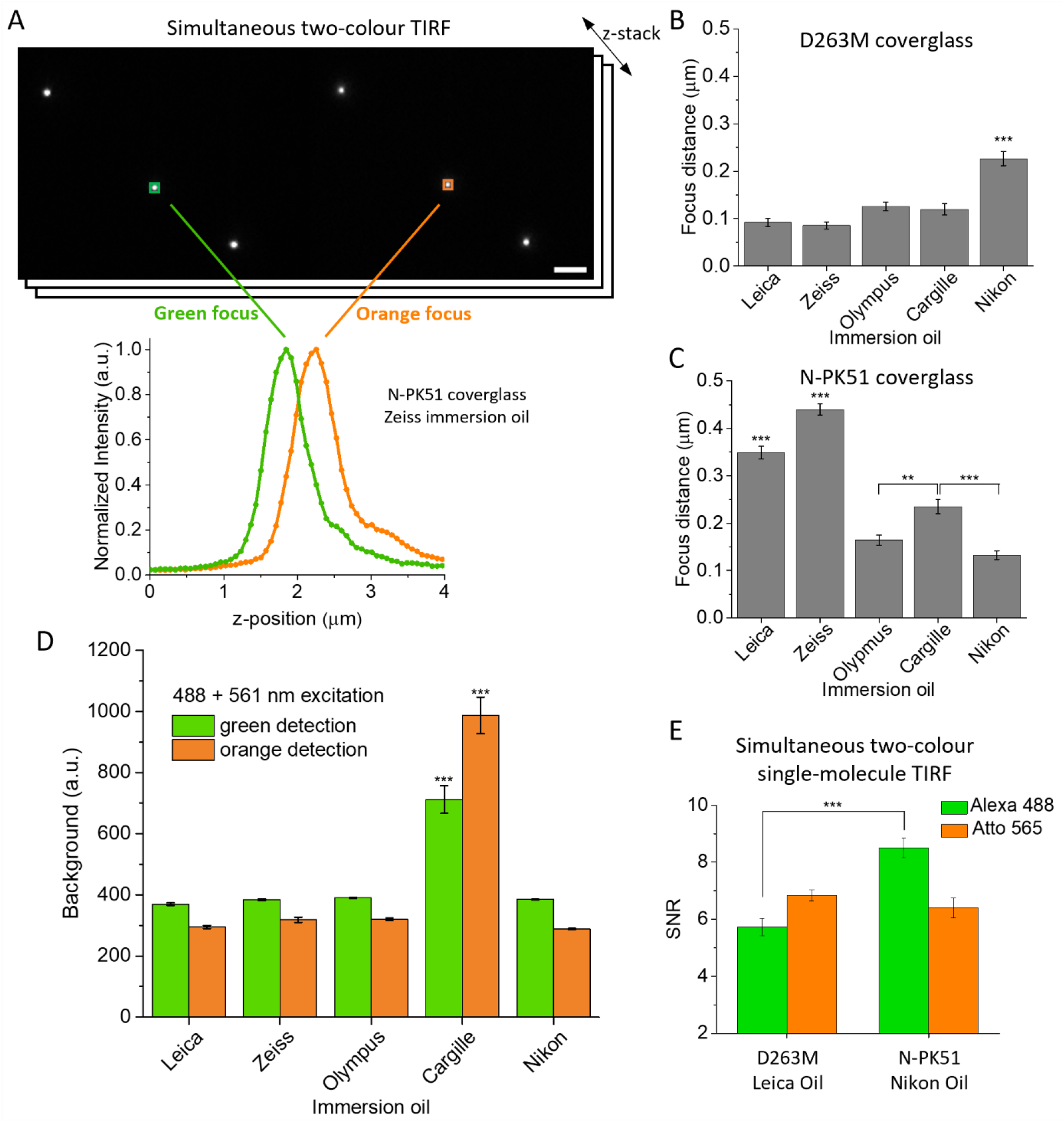
Effects of immersion oil on chromatic aberrations and background. A) Chromatic aberration is evident from the graphs of intensity z-profiles of single fluorescent beads able to emit in different wavelength bands, imaged simultaneously in the two channels (green and orange graphs for emissions in the green and orange channels). On top, example of images taken at 2 µm. Scale bar: 3 µm. B) Distances between the focus z-positions (the one corresponding to the maximum intensities in panel A) of the two channels using different immersion oils and D263M cover glass. C) Same as B) with N-PK51 cover glass. Data are mean ± SEM, obtained from profiles of 30 beads in each condition. *** P < 0.001, ** P < 0.01, 1-way ANOVA, Bonferroni multiple comparisons. D) Background detected in TIRF under simultaneous 488+565 nm excitation using different immersion oils (measurements performed with D263M cover glass). Data are mean ± SEM, obtained from 7 different fields of view. *** P < 0.001, 1-way ANOVA, Bonferroni multiple comparisons. Tests performed separately on the green and orange results sets. E) SNR for single molecules of Alexa 488 and Atto 565 adherent on the cover glass in a TIRF configuration with simultaneous excitation and detection. Measurements were performed using the D263M cover glass with Leica immersion oil (left) and the N-PK51 cover glass with Nikon immersion oil (right). Data are mean ± SEM, obtained from 5 movies in different fields of view, with 2000-20000 spots analyzed in each movie. *** P < 0.001, Student’s test. Tests performed separately on the green and orange couples of measurements.

We also compared the background detected in the system using various immersion oils developed for fluorescence microscopy. Based on our results, we excluded Cargille LDF, while the others showed comparable behaviors (Figure 3D). With the identified best conditions in our system (collar position and immersion oil) for each cover glass, we measured the SNR in a real two-color TIRF configuration with simultaneous double excitation and detection. On dyes adhered to the cover glasses, we found no significant differences for the 565-dye; instead, we found a significant improvement for the 488-dye using the N-PK51 cover glass (Figure 3E).

### 2.4 Cell autofluorescence

Another known source of autofluorescence is the cell itself [21]. We measured cellular autofluorescence with the chosen excitation-detection couples for the two-color configuration based on 488 and 565 dyes, i.e. 488 nm excitation with green and orange detection and 561 nm excitation with orange detection. Moreover, we performed the measurement using 635 nm excitation with far-red detection in order to check if the impact of cellular autofluorescence can affect our conclusions as reported above, since this is the single-channel configuration typically chosen in fluorescence microscopy. We performed measurements at different TIRF penetration depths (Figure 4). Considering the longest penetration depths, it is clear that the excitation at 488 nm induced higher emissions than the excitations at 565 nm and 635 nm. The contributions in the green or orange range under 488 nm excitation were similar. However, for penetrations of 130 nm and lower, the contribution to background due to cell autofluorescence did not exceed 5% of the measured background even in the case of excitation at 488 nm, and it reached 2 % for 100-nm penetration-depth. 100 nm is the typical penetration depth used for single-molecule imaging and tracking of cell-surface proteins [25]. It was observed that cell fluorescence arises from intracellular components and it is most intense in discrete cytoplasmic vesicles [21]. Indeed, we found higher contributions at deeper penetration and usually observed that autofluorescence did not have a homogenous pattern in the cell area, but was concentrated in some regions (see Figure 4B).

**Figure 4.**
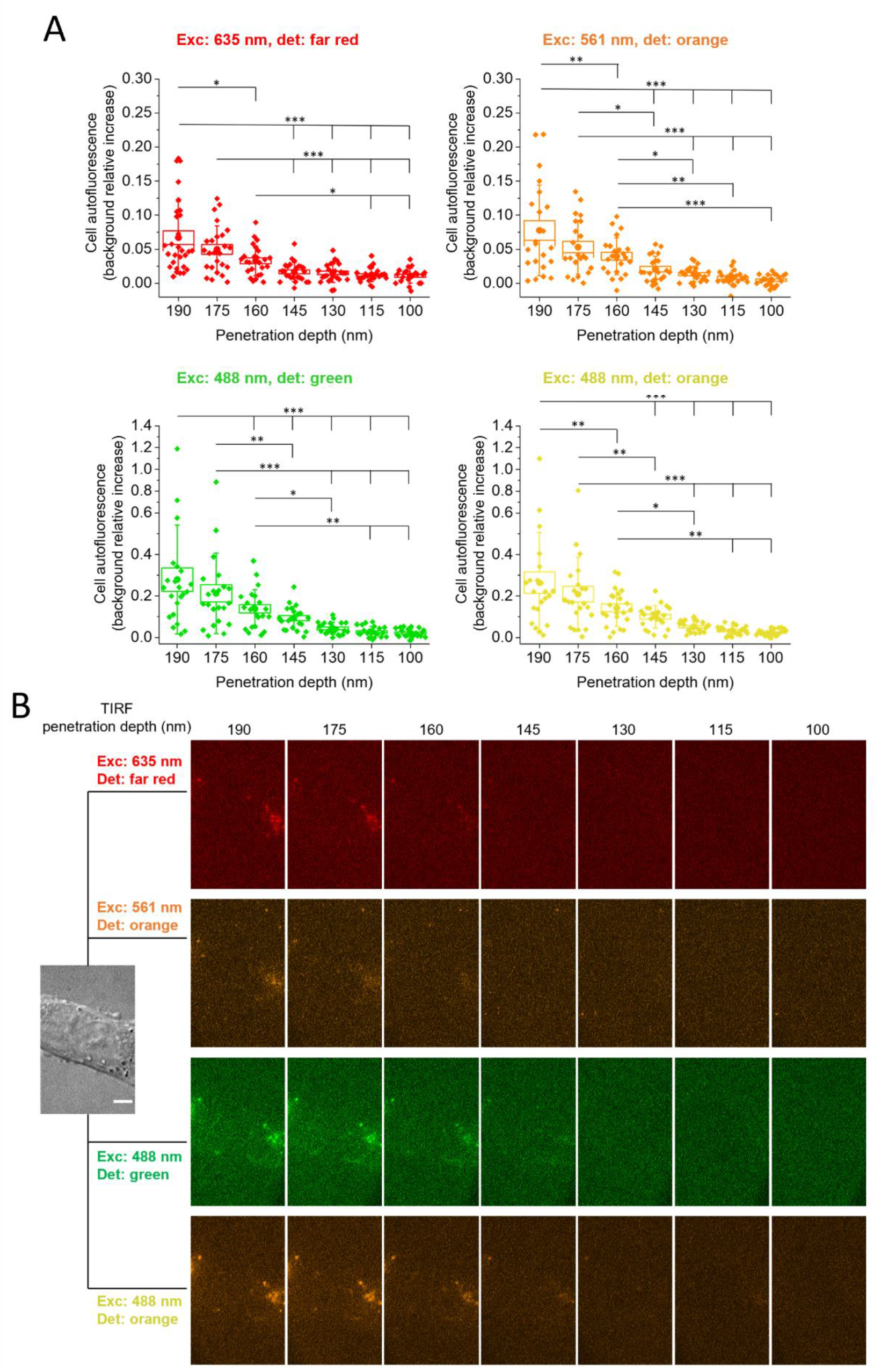
Cell autofluorescence in TIRF. A) Cell autofluorescence measured on SK-N-BE(2) cells for different excitations (Exc) and detections (det) wavelengths, using different TIRF penetration depths. The Y axis reports the relative increase of background due to cells autofluorescence (i.e., the ratio between the background due to cell autofluorescence and the background due to all other sources, see also Supplementary Methods). * P < 0.05, ** P < 0.01,*** P < 0.001, 1-way ANOVA, repeated measurements, Tukey multiple comparisons. Box: SEM, whiskers: standard deviation, spots: single cells from two independent repetitions. B) representative images of the experiment: a cell (DIC image on the left) is visualized with the different excitation-detection combinations at different TIRF penetration depths.

### 2.5 Cell medium autofluorescence

Highly sensitive live-cell imaging calls for the optimization of the cell-imaging medium. Indeed, traditional growth media typically induce autofluorescence signals that negatively impact the available SNR [26]. On the other side, imaging in balanced salt solutions lead to poor cell health. Typically, the higher growth media emissions are induced by UV, violet (405 nm) or blue (488 nm) excitation, while absorbance above 600 nm is usually not significant [26]. A correlation between the vitamin content and media autofluorescence induced at excitations lower than 500 nm, including 488 nm was observed [26,27]. Low-fluorescence media formulations with optimized vitamins (e.g. riboflavin) were developed [26,27]. Among these, FluoroBrite™ DMEM was tested for long-term live-cell imaging and cell culture maintenance, even on vitamin-sensitive and nutrient-sensitive cell lines [27].

In order to monitor the impact of cell media fluorescence, we compared emissions from saline buffers and the cell media RPMI, DMEM, DMEM/F12, FluoroBrite DMEM (all red-phenol-free). We observed significantly higher emission with 488 nm excitation from cell media RPMI, DMEM and DMEM/F12 with respect to PBS, in both green and orange regions (Figure 5). An element likely involved in this behavior is riboflavin since, under excitation at 488 nm, it emits between 500 and 650 (green/red) with a main peak in the green region at 535nm [28,29]. Riboflavin is present in all three media; however, in DMEM it has a concentration of 0.4 mg/L, twice the value in the other two media (information available from the producer Thermofisher Scientific). This may explain the higher fluorescence detected with DMEM. Importantly, FluoroBrite™ DMEM fluorescence was not significantly different from PBS or from a solution of HEPES and salts in our tests (Figure 5; see Supplementary Methods for the detailed composition); moreover, we did not find any significant difference when adding some common supplements (Figure S4).

**Figure 5.**
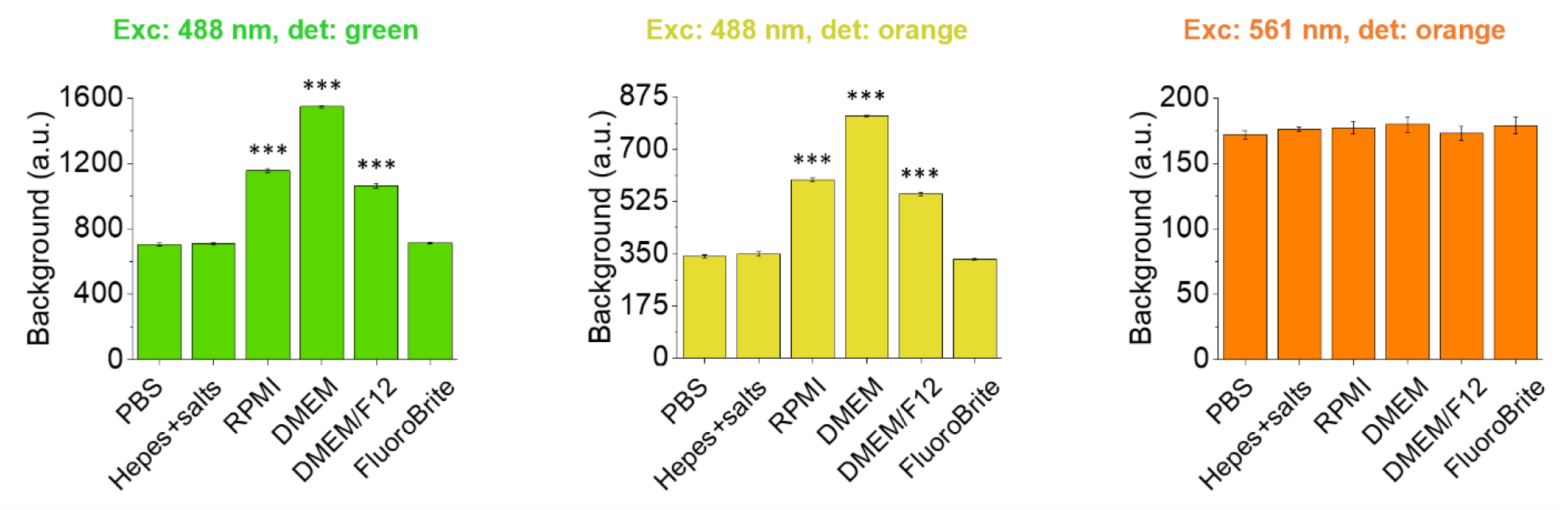
Autofluorescence from cell media. Background detected in TIRF using different cell media. Results are mean ± SEM, obtained from 6-16 different fields of view from two independent repetitions. *** P < 0.001, 1-way ANOVA, Bonferroni multiple comparisons.

### 2.6 Photostability improvement for two-color imaging of 488 and 565 channels

After SNR optimization, we addressed photobleaching, a severe limitation for single molecule tracking. First, we compared the photobleaching lifetimes (PBLTs) of the most popular 488-dyes used in single-molecule imaging (i.e., Alexa 488, Atto 488, Abberior STAR 488) (Figure S5 A). We found significant differences, with Atto 488 being the most photostable. The brightness of Atto 488 is the highest as well (even if not by much; see Table S1). Atto 565 was more photostable than the 488-dyes in our experimental conditions. Therefore, we selected Atto 488 and Atto 565 for the following measurements (Figure S5 B).

Various antifading agents were proposed for use in cell cultures, among which ascorbic acid (AA), n-propyl gallate (NPG), and Trolox (TX) [30,31]. Higher efficacy of TX was reported, in some cases, when it was partially oxidized to form a small percentage of Trolox-quinone (TQ), acting therefore as a reducing-plus-oxidizing system (ROXS) [23,31]. Since this effect depends on the considered dyes [23], we looked for the optimal TQ percentage for the two dyes of interest. We first monitored Trolox oxidation under UV illumination [31] (Figure S6 A, B). In absorbance measurements of a fresh Trolox solution, we observed its characteristic absorbance peak at about 290 nm; using different UV-excitation times, we observed the formation of a new peak at about 270 nm due to the TQ with the expected isosbestic point at about 285 nm [31]. For each UV illumination time, we extracted the corresponding percentage of formed quinone (Figure S6 B) and the PBLT of the two dyes observed in TIRF measurements in the presence of the corresponding solution (Figure S6 C). In both cases, we observed that the best condition is the use of Trolox without oxidation, with an approximately linear decrease in PBLT as the quinone concentration increased. The decrease of photobleaching lifetime cannot be ascribed simply to a lower effect of Trolox due to its lower concentration, rather it seems that the TQ increases the photobleaching of the considered dyes. It was reported that the ROXS approach tends to give greater improvements for far-red-emitting dyes (such as Atto 655, Atto 647N, Cy5 [31,32]) and little or no improvements on fluorophores excited at shorter wavelengths [33,34]. This was attributed to an increasing influence of higher-excited states in photobleaching pathways over the triplet state (suppressed by ROXS) as the excitation energy increases. Furthermore, higher energy excitations may enhance the creation and impact of reactive oxygen species (ROS). TX was used in non-oxidized form in all the following measurements.

The best concentrations of TX, AA and NPG as antibleaching agents are not univocally determined in the literature. TX was used at 1-2 mM [23,32,35], AA was used in mM concentrations (1-10 mM) but also in the *μ*M range [30,34,36], NPG was used at concentrations ranging from few *μ*M to 10 mM [30,37]. Therefore, we measured the effect of different concentrations of the three reagents separately on the two dyes Atto 565 and Atto 488 (Figure 6 and S5 B). We extracted the intensity-decay profiles that were fitted by an exponential function to extract the PBLT (see Supplementary Methods). We found that each of the reagents had a positive effect on the PBLT for both dyes. However, there are several differences in the observed effects. Comparing the plateau PBLTs, TX was the most effective additive for Atto 488, while AA and NPG had similar effects; NPG was the most effective and AA was the least effective on Atto 565. Moreover, for Atto 488, TX and AA had concentrations of a few mM at the plateau, while NPG required just a few *μ*M. Similarly, Aitken et al. noted that 100 *μ*M NPG (used in their work at lower concentration just due to its lower solubility) caused the same improvement on photobleaching as 10 mM AA on Alexa 488 and Cy3 [38]. Also for Atto 565 a few *μ*M NPG allowed reaching the plateau; TX and AA required concentrations were lower (about 50 *μ*M) than for Atto 488.

**Figure 6.**
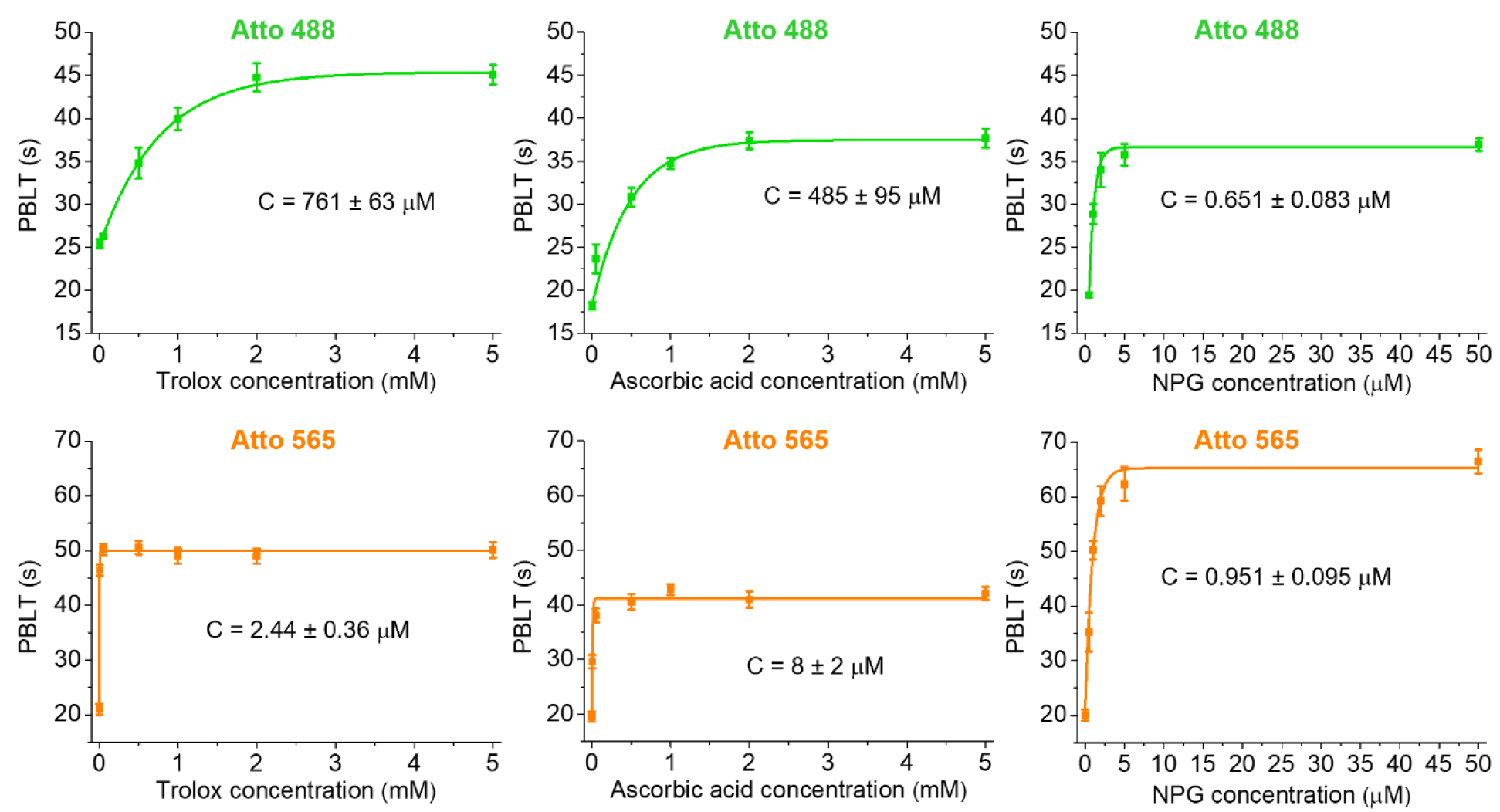
Effects of different reagents on photobleaching. Photobleaching lifetimes (PBLTs) of Atto 488 (top, green) and Atto 565 (bottom, orange) in the presence of different concentrations of Trolox, Ascorbic Acid and n-propyl gallate (NPG). Each lifetime value was obtained by fitting the temporal decay of intensity observed in those conditions (see Figure S5 and Supplementary Methods). PBLTs versus reagent concentrations were here fitted with a monoexponential function y = A exp (-x/C) + y0. Obtained C values are reported on each graph. Lifetime data were acquired in TIRF from 10-30 different fields of view from samples of 2-4 independent repetitions. Results are mean ± SEM.

We then tested the effects of combinations of different reagents (Figure 7). We first fixed NPG concentration (at 2 *μ*M) and compared the effects of mixing it with AA, TX and a combination of TX and AA (by using 5mM for both TX and AA, where their plateaus were already reached for both dyes) (Figure 7A). On Atto 488, we measured a longer PBLT with a mix of TX and NPG than with AA and NPG; we did not find further improvement by using the three reagents together. For Atto 565, we did not find significant differences between the three situations. We discarded therefore the combination of NPG with AA and of the three reagents together and compared a mix of TX plus AA with a mix of TX plus NPG (by fixing TX concentration at 5mM and using 5mM for AA and 5 *μ*M for NPG, concentrations at which the plateaus were already reached); for both dyes, we observed a significantly longer PBLT when using the combination of TX and NPG (Figure 7B). The comparisons reported in Figure S7 show that using 5 *μ*M NPG is significantly better than using 2 *μ*M NPG, at least for Atto 488 (and it is not worse for Atto 561).

**Figure 7.**
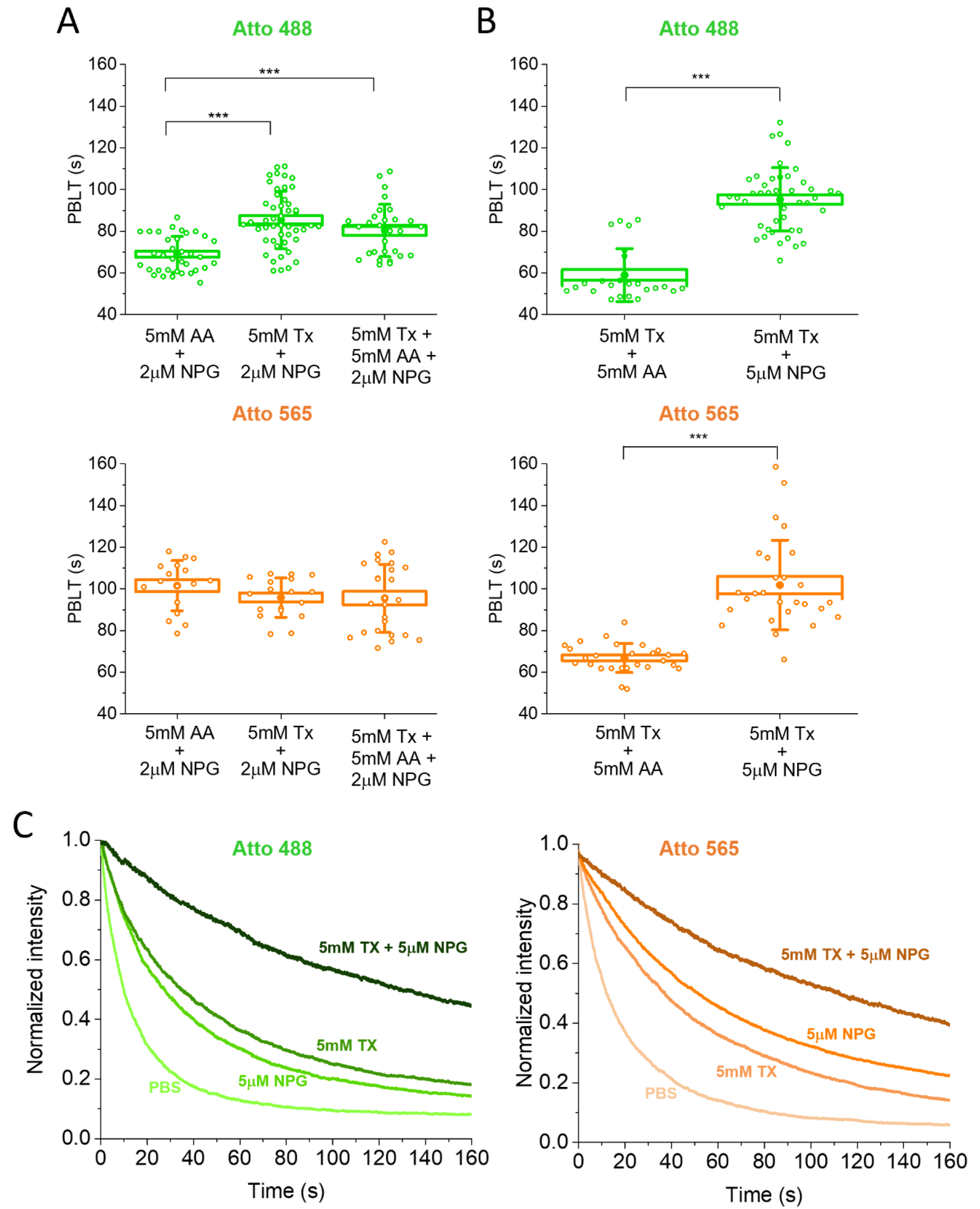
Reduction of photobleaching with mixed reagents. A) Atto 488 (top, green) and Atto 565 (bottom, orange) photobleaching lifetimes (PBLTs) observed adding a mixture of ascorbic acid (AA), Trolox (TX) or both with a fixed concentration of n-propyl gallate (NPG). *** P < 0.001, 1-way ANOVA, Bonferroni multiple comparisons. Box: SEM; whisker: SD; dots: individual fields of view, from 2-4 independent repetitions. B) As in A, for a mixture of TX and AA and a mixture of TX and NPG. *** P < 0.001, Student’s test. C) Examples of temporal decay of intensity observed in TIRF under different medium compositions for Atto 488 (left, green) and Atto 565 (right, orange).

These data indicated that in all cases and for both dyes, the observed PBLTs were higher with mixed reagents than with single reagents at their plateau, likely thanks to a simultaneous suppression of different bleaching pathways; importantly, the best combination found (TX plus NPG) was the most effective for both dyes (Figure 7). A recent study that searched for the best photobleaching conditions on different dyes by comparing TX, ROXS or deoxygenation plus ROXS, on Atto 488 obtained a final improvement in PBLT of 4.4 times (Atto 565 was not considered in the study) [23]. Here, on Atto 488 we achieved an improvement of about 6 times (14 to 95 ms) with the combination of TX+NPG. Another work considered Atto 565 comparing: oxygen depletion plus AA or MV (methyl viologen) or ROXS (based on AA plus MV), TX plus MV, TX alone; the authors could not find a solution significantly better than TX alone [32]. Here we showed that a better solution is its combination with AA and especially with NPG.

### 2.7 Two-color single-particle tracking on fast diffusing p75NTR receptor

The examined improvements are particularly important to study interactions of single fast-moving molecules. In order to test if the best-identified experimental conditions determined above are sufficient, we performed simultaneous two-color single-particle tracking (SPT) on the p75^NTR^ membrane receptor in living cells.

We used labelling through Sfp phosphopantetheinyl transferase (Sfp). This strategy allows a covalent conjugation of the dye to a serine residue of a peptide tag inserted in the receptor sequence resulting in a controlled 1:1 stoichiometry between dye and receptor [39–41]. We used S6-tagged p75^NTR^, as previously used in single-color SPT studies on this molecule [7,42].

First, we performed single-color experiments on SK-N-BE (2) cells cultured in petri dishes for microscopy with thin bottom cover glass made of standard D263M and of N-PK51 materials. We labelled the receptors with Atto 565. We confirmed that experiments on living cells using the new type of cover glass are feasible. We compared the diffusion coefficient distribution in the two cases and we obtained the same results on the two cover glasses (Figure S8). We then performed simultaneous two-color SPT by labelling p75^NTR^ receptors with a mixture of Atto 565 and Atto 488. First, we carried out the experiment on D263M cover glass (Fig 8 A, C, Supplementary Videos 1, 2, 3). We measured the diffusion coefficient distribution, comparing it with the one obtained in the single-color case. We observed that in the two-color configuration only the 565-channel produced results comparable with the single-color experiment while the 488-channel produced a distribution shifted towards higher diffusivities (Figure 8 C). We also performed the two color SPT experiment with a different receptor, i.e. TrkA, and also in this case, we observed a shift towards higher D for its distribution when estimated on the 488-channel, compared to the 565-channel (Figure S9). We then performed two-color SPT on p75 receptors using N-PK51 cover glasses (Fig 8 B, C; Supplementary Videos 4,5,6). In this case, we obtained the same distribution of the diffusion coefficients in both the 565 and 488 channels, the same obtained in the single-color case. Thus, the shift for the 488-channel observed using the D263M cover glass was likely due to the low SNR in that channel, which causes significant tracking errors due to high percentages of missed detections. To confirm this hypothesis, we simulated movies of single molecules diffusing with a given diffusivity, mimicking conditions (signal intensity, background, noise) similar to those observed in experiments on D263M cover glass (Figure S10, Table S4); we then performed the SPT analysis on the simulated movies. We observed the same shift of the diffusivity histogram peak for the situation corresponding to the 488 channel (Figure S10). There are some differences between experiment and simulation; in particular, experimental distributions are broader, especially for the 565-channel case. This can be explained by the fact that in the simulation we did not consider some effects present in cells, like an intrinsic spread in D values (caused by membrane areas with different compositions or by interactions with membrane structures and other molecules) or the impact of immobile dye molecules nonspecifically adsorbed on the cover glass. While the experimental distributions of 565 channel were broader mainly due to these heterogeneities, the distribution width in the 488 channel appears to be dominated by tracking errors and thus is similar between experimental and simulated movies. The impact of immobile dyes causes a shoulder in the left tail of the distribution, which is higher for Atto 565 than for Atto 488; this is caused by a stronger non-specific adhesion for the first dye, as observable in our results and reported in the literature [43,44].

**Figure 8.**
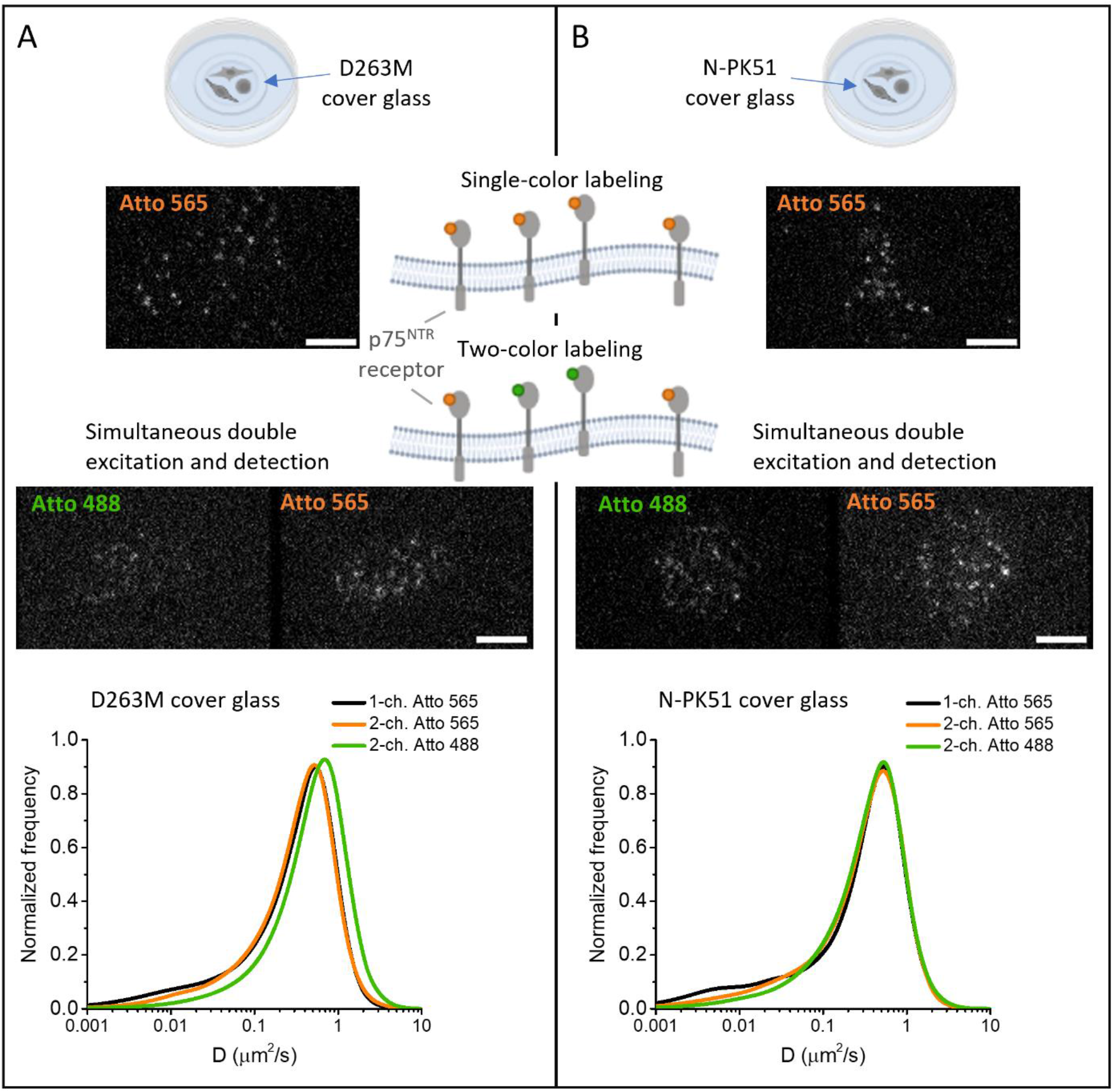
Single particle tracking of p75 receptors. One-color and two-color SPT were performed on SK-N-BE cells grown on D263M (A) and N-PK51 (B) cover glasses. p75 receptors were labeled with Atto 565 for one-color experiments or with a mix of Atto 565 and Atto 488 for two-color experiments. TIRF representative images from the acquired movies are shown (scale bar: 5*μ*M). On the bottom: distributions of diffusion coefficient (D) extracted with single particle tracking analysis on one-color acquisitions (1-ch. Atto 565, black curves), simultaneous two-colour acquisitions (2-ch.; orange curves for Atto 565 and green curves for Atto 488). D263M cover glass, 1-ch Atto 565: 4086 tracks. D263M cover glass, 2-ch Atto 565: 2501 tracks. D263M cover glass, 2-ch Atto 488: 2119 tracks. N-PK51 cover glass, 1-ch Atto 565: 2127 tracks. N-PK51 cover glass, 2-ch Atto 565: 1828 tracks. N-PK51 cover glass, 2-ch Atto 488: 2086 tracks. Trajectories were obtained from 6-8 different cells from two independent repetitions in each case.

In conclusion, with standard cover glasses it was not possible to obtain reliable results in both channels in two-color SPT even minimizing the autofluorescence of medium, cells and oil. The challenge was overcome by using the N-PK51 cover glasses, which yielded higher SNR in the 488-channel and reliable results in two-color SPTs, even with fast receptors such as p75^NTR^.

## 3. Discussion

The development of ever more sensitive single-molecule microscopy applications requires continuous efforts to overcome limitations imposed by the experimental setups, from microscopy components to fluorescent probes [45– 49].

In this work, we addressed two essential requirements for quantitative single-molecule imaging and tracking, i.e. signal-to-noise ratio and photostability, achieving a simultaneous two-color TIRF configuration able to visualize fast-diffusing membrane receptors labelled with minimally-invasive dyes in living cells.

Reaching this goal required understanding and dealing with different background sources within the setup, which significantly affect the SNR of single molecules. In particular, we demonstrated that multicolor applications are mostly hindered by the background resulting from the cover glass, whose standard material is borosilicate glass. Its emission in red and far-red regions under excitation at shorter wavelengths strongly limits the use of those spectral bands for detecting single molecules in multicolor configurations. We searched for different possible materials to be used as cover glasses in TIRF systems that, especially considering commercial ones, have strict requirements on the properties of the cover glass, i.e. thickness of 0.17 mm and refractive index of 1.52. This hinders the use of quartz and fused silica which have much lower autofluorescence than glass [50,51] but a refractive index of 1.46 that causes aberrations in setups designed for a 1.52 value.

After extensive searching, we selected Schott N-PK51 glass for making custom cover glasses (See Supplementary note and Figure S2). Our tests showed a significantly reduced autofluorescence in the green region under 488 nm excitation compared to standard cover glasses. As a consequence of this, the material allowed a significant improvement of the SNR for simultaneous two-channel imaging based on “488” and “565” dyes, which we identified as the best couple for this kind of experiments. The slightly different optical properties of N-PK51 compared to the standard material can be addressed even in a commercial system, thanks to small corrections introduced by the objective collar and the immersion oil.

For live-cell experiments, two other background sources were considered, namely cells and cell medium. Most cell autofluorescence is excited at wavelengths below 450 nm [52]. However, in various cell types, the 488 nm wavelength excites flavins which emit in the range 500-650 nm with a peak at about 550 nm [29,53]. The emission in orange and red regions excited at 561 nm or 635 nm can instead be associated with porphyrins [46]. The latter have a main absorption peak around 400 nm (Soret band) plus minor bands between 500 and 700 nm (Q bands) and emit in the orange-red region (between 600 and 700 nm) [53,54]. Since this fluorescence emission is usually lower than that excited at 488 nm, single-color fluorescence measurements are often performed using far-red-emitting dyes. However, we demonstrated that by selecting suitable TIRF penetration depths for SPT on the cell membrane, the contribution of cells to the background can be less than 2% under 488 nm excitation. Furthermore, we identified a cell medium (FluoroBrite™ DMEM) that allowed us to reduce background emissions to levels comparable with those of PBS; this medium ensures cell health and avoids the use of saline buffers alone, which lack essential elements for cells. Thus, we could confirm that the most limiting factor for TIRF-based multicolor single-molecule imaging is the background from the cover glass. So, the choice of channels should be based mainly on the characteristics of this emission rather than on those of cells and cell medium (as long as these are properly optimized) and therefore we could confirm the selection of the 488-565 dye pair.

Alongside SNR, the second crucial point in quantitative single-molecule microscopy, especially when using small and minimally invasive organic dyes, is photobleaching, which limits the number of photons emitted before photodestruction of the marker. Studies on photobleaching have revealed the complexity of the phenomenon, still poorly understood and stemming from different, dye-dependent mechanisms, including interactions with reactive oxygen species (ROS), and reactions of excited states (especially triplet ones) of the dye [22,30]. Several methods were used to reduce photobleaching, from deoxygenation to the addition of triplet state quenchers or reducing and oxidizing reagents [55]. We focused on three reagents with potential antibleaching properties: Trolox (TX, a free radical scavenger, an antioxidant and possibly a triplet state quencher), ascorbic acid (AA, well-known antioxidants with a possible scavenging activity towards some free radicals) and n-propyl gallate (NPG, antioxidant and scavenger for some ROS) [30,31,55,56]. In general, the action of each reagent and the best solution against photobleaching strongly depend on the dye being considered: some strategies effective on some dyes may not be effective or have adverse effects on others [23,32,35]. For this reason, multicolor studies have the additional challenge of finding a solution that works well for multiple dyes simultaneously. We achieved this objective on the two dyes of interest Atto 488 and Atto 565. Both showed less photobleaching when adding TX, AA or NPG. For both dyes we found the most effective solution with a combination of TX and NPG.

We applied the developed tools to perform two-color SPT experiments in live cells.

We selected the labelling based on Sfp phosphopantetheinyl transferase as this minimizes perturbations thanks to a controlled 1:1 stoichiometry between dye and receptor and the insertion of only a very short tag (12 residues) in the protein sequence [39–42]. We chose the p75^NTR^ receptor for the labeling with two small organic dyes to perform two-color imaging. This choice was driven by the observed alteration of its dynamics upon labeling with Qdots with respect to small organic dyes [7]. More importantly, this is a particularly fast receptor and, as we also showed here, higher diffusivities create worse SNR conditions. We demonstrated that the presented methods allow reliable single-molecule tracking simultaneously in two channels even on challenging receptors, unlike standard TIRF conditions, effectively extending the potential of TIRF-based SPT.

## 4. Materials and methods

### Microscopy setup

Microscopy experiments were carried out with a Leica DM6000 inverted microscope (Leica Microsystems) equipped with an epifluorescence module, DIC in transmission, TIRF-AM module, HCX PL APO 100.0X oil-immersion objective (NA1.47), EM-CCD camera (iXon Ultra 897, Andor), four laser lines (405, 488, 561 and 635 nm), incubator chamber to maintain 37°C and 5% CO2 conditions for live cell imaging.

An external laser combiner (iFLEX-adder, QiOptiq) with kineFLEX polarization maintaining fibers (QiOptiq) and kineMATIX fiber coupler (QiOptiq) was used with a 488 nm supplementary laser (iFLEX-iRIS, Qioptiq) for simultaneous two-color excitation. The power of the additional 488 nm laser was tuned by a DAC. To detect two channels at the same time, a Dual View (Optical Insights DV-CC) was placed in front of the EM-CCD camera, so that emitted light was split and collected in the two halves of the camera. We included an Optomask adjustable field mask (OPTMSK-L, Andor) between the microscope output and the Dual View to limit the illuminated area of the camera and increase the frame rate.

We used the fluorescence cube Leica Quad ET TIRF MC for Laser lines 405, 488, 561, and 635 nm (except when differently stated). In the Dual View, two filter sets were used: i) dichroic beam splitter T600lpxr, with filters Chroma ET525-50 and r647lp, for detection of green-emitting dyes and far-red-emitting dyes respectively; ii) dichroic beam splitter 565dcxr, with filters Chroma ET525-50 and Semrock FF01-600/52, for detection of green-emitting dyes and orange-emitting dyes respectively. For single-color comparisons between Atto 565 and Abberior STAR 635p we also used the Cy3 and Cy5 cube as specified in the SNR measurements section.

### SNR measurements

We measured signal-to-noise ratio (SNR) on fluorophores adhered to the bottom of WillCo® dishes with their original D 263® M cover glass or with custom cover glasses made of Schott N-PK51 glass (Figure 1 B, D, 2 B, C, D, 3E, S3). We also performed SNR measurements on fluorophores that labeled membrane receptors TrkA and p75 in living cells (Figure 1B and S1, see also Supplementary Methods).

In all cases, samples were prepared with fluorophore densities low enough to distinguish single molecules. For measurements on adhered fluorophores, a solution of them in PBS (typical concentration of about 1nM) was deposited on the cover glass for about 10 minutes. After extensive washing to remove fluorophores remained in suspension or weakly adhering, the dish was filled with PBS and the sample was immediately observed with the TIRF microscope.

For single-color comparisons between Atto 565 and Abberior STAR 635p (Figure 1B), we used 561 nm excitation with a Cy3 cube and 635 nm excitation with a Cy5 cube, respectively, with both laser power set to 3 mW at the objective.

For single-or double-color measurements on Alexa 488 and Atto 565, we used single or double excitation at 488 (3.5 mW) and 561 nm (3 mW), with cube and filters as specified in the Microscopy setup section.

For preliminary evaluations of the SNR achievable in two-color configurations with different pairs of channels (without needing setups for simultaneous excitation and detection for all possible pairs), we exploited the Dual View system detailed above considering one window at a time under single excitations. We performed single-color acquisitions on fluorophores for each one of the considered channels; then, for each possible pair of channels, we measured the background resulting upon excitation at the shorter wavelength in the longer wavelength detection channel. After subtracting the offset, we added this background to the images acquired under the single excitation. This allowed to estimate the SNR of the channel at longer wavelengths upon double excitations. In this way, we compared three couples of channels with excitations at 488+561 nm, 561+635 nm and 488+635 nm excitation (Fig 1D).

We acquired 100-frame time series with 40 ms integration time in different fields of view for each sample.

Analysis were performed with TrackMate 6.0.1 plugin in ImageJ [57]. After detection and tracking, we considered spots in tracks lasting at least four frames to exclude false detections. For measurements in live cells, we also applied filters on the displacement and velocity of tracks to select only dyes labelling moving receptors and exclude the ones non-specifically adhered to the glass. For each field of view, the SNR was then calculated by subtracting the background measured in the same field of view in dye-free areas from the mean spot intensity (calculated by TrackMate using a fixed spot radius) and dividing the result by the standard deviation of that background. For in-cell evaluation, the background was evaluated in areas with non-transfected cells.

### Single-particle tracking

After labelling, cells were immediately imaged by TIRF microscopy. We selected a constant region of interest (ROI) containing at least part of the basal membrane of each cell; ROI were 318 × 236 pixels (50.88 × 37.76 μm, pixel size 0.16 μm) for two-channel acquisitions and 159 × 236 pixels for one-channel acquisitions. We acquired 250-frame time series with an integration time of 40 ms, corresponding to a frame time of 56 ms. Penetration depth was set to 110 for the 561 nm internal excitation, corresponding to a depth of about 96 nm for the 488 nm external one in the case of a simultaneous two-color acquisition.

For the tracking analysis, we first performed some pre-processing steps with the ImageJ software: i) in the case of two-channel acquisition, we split the movie into two movies, one for each channel; ii) for each movie we created a maximum intensity projection to identify the contours of the cell membrane; iii) we applied a mask to the movie using the identified contours to exclude the spots outside the cell due to the fluorophores adhered in a non-specific way to the cover glass, analogously as in [7,58]. The masked movies were then processed with u-track software version 2.2.1 [59] in MATLAB to obtain single-particle trajectories.

For the calculation of distribution coefficients (Figure 8, S1, S8, S9), we selected tracks including at least 6 detected spots and lasting at least 9 frames in total (including gaps). We divided subtrajectories created by merge and split events, as described in [7]. On the obtained subtrajectories, we calculated the diffusion coefficient D from the first two points of the Mean Square Displacement (MSD) function. The D distributions were determined considering uncertainties on D and weighting each trajectory with the number of frames where its spots have been detected, as in [58]. Distribution integrals were normalized to 1.

More details on material and methods appear in Supplementary Methods.

## Acknowledgements

We acknowledge Laura Marchetti and Fulvio Bonsignore for developing the initial versions of some of the constructs used in this work. We acknowledge Filippo Fabbri for measurements with the Renishaw micro-Raman system.

This research received funding from Scuola Normale Superiore (SNS16C_B_LUIN, SNS_RB_LUIN, SNS19_A_LUIN), from Fondazione Pisa (project Nanotechnology for tumor molecular fingerprinting and early diagnosis, RST 148/16), and from the European Union Next-GenerationEU through the PIANO NAZIONALE DI RIPRESA E RESILIENZA (PNRR - MISSIONE 4 COMPONENTE 2) within the National Quantum Science and Technology Institute (NQSTI - INVESTIMENTO 1.3; PE_00000023) and the Tuscany Health Ecosystem (THE - INVESTIMENTO 1.5; ECS_00000017).

## Author contributions

Conceptualization, C.S.S. and S.L.; Methodology, C.S.S., S.L., R.A., A.M.; Validation, C.S.S.; Software, C.S.S., S.L.; Formal Analysis, C.S.S.; Investigation, C.S.S.; Resources, F.B., S.L.; Data Curation, C.S.S.; Writing – Original Draft, C.S.S., S.L.; Writing – Review & Editing, C.S.S., R.A., A.M., F.B., S.L.; Visualization, C.S.S., S.L.; Project Administration, S.L.; Supervision, S.L., F.B.; Funding Acquisition, F.B., S.L.

