## Supplementary Information for "Optimized two-color single-molecule tracking of fast-diffusing membrane receptors"

### **Supplementary Note**

Information on the characterization of optical glasses autofluorescence is still very limited in the literature. The very few reported comparisons [1–3] include a very limited number of glasses compared to the existing ones; D263M, the typical cover glass material, is never included; data are mainly obtained under UV-excitation, and even more limited information exists for excitation in the visible range; reported data are sparse and difficult to compare with each other, in particular because there is no well-defined standard or reference; glass datasheets do not include information about this phenomenon and it is not possible to compare glasses from different companies. We only found recent attempts by the Schott company to provide some comparisons of the autofluorescence of its glasses. The Abbe diagram of Schott glasses shows that the classes with refractive index closest to 1.52 (the standard value for cover glasses) are BK, K, KF and PK classes (Fig. S2). A technical note by Schott reports that PK glasses (51 and 52A) have an integral fluorescence under 365, 405, 488 and 532 nm excitation that tends to be low (relative to the subset of glasses considered in the measurement) and lower than BK7 (a borosilicate) [4]. Their integral fluorescence at 642 nm excitation instead is among the highest and is higher than BK7. K class glasses (K5 and K10) have higher integral fluorescence than BK7 for 365, 488, 532 nm; for 405 nm excitation, K5 has lower integral fluorescence than BK7; for 642 nm excitation, K5 has higher integral fluorescence than BK7 and lower than PK51 and PK52A [4]. There are no further data for the other glasses of the PK, BK and K classes and none for KF class.

For none of the classes of interest, an emission spectrum is provided together with information on the integral fluorescence. The integral fluorescence was measured in the ranges: 370-700 nm for 365 nm excitation, 535-800 nm for 488 nm excitation, 580-850 nm for 532 nm excitation; for 405 and 642 nm excitation, it is stated that limits were defined accordingly [4]. However, the shape of the spectra is important to know in which bands the main emissions are, especially to set-up multichannel configurations.

Within the PK class, for excitations at 488, 532, 642 nm, PK51 has a lower autofluorescence than PK52A [4].

From this scattered data, we selected PK51 glass to make customized cover glasses to be tested in our system.

### **Supplementary Methods**

#### *Synthesis of CoA-fluorophores*

Alexa 488 maleimide was obtained from Molecular Probes (Invitrogen, Eugene, OR); Atto 488 maleimide and Atto 565 maleimide were obtained from ATTO-TEC (Siegen, Germany); Abberior STAR 635P maleimide was obtained from Abberior (Goettingen, Germany). Coenzyme A trilithium salt was obtained from Sigma-Aldrich. All solvents (ultrapure grade) were obtained from Sigma-Aldrich.

Chromatographic analyses and purifications were performed using a Phenomenex Kinetex EVO C-18 150 x 3.00 mm column on a Dionex Ultimate 3000 high-performance liquid chromatography system (UHPLC) equipped with a photodiode array (PDA) detector. For analyses, the HPLC was interfaced with an ABSciex API 3200 Q-Trap mass spectrometer (MS). Concentrations of CoA-dyes after purification were determined using a UV/vis spectrophotometer (Jasco V-550).

For the synthesis of all CoA-fluorophores, 0.412 μmol of CoA were first dissolved in 100 μL of a Tris(2-carboxyethyl)phosphine (TCEP) solution (10 mM in PBS) and left stirring 1 h at 37°C. 0.375 μmol of the fluorophore were dissolved in 40 μL of DMF. CoA solution was cooled to room temperature, the fluorophore was added and the reaction was stirred for 2-4 h at 37°C. Then, CoA-conjugated fluorophores were purified by the described HPLC system, using Ammonium acetate buffer 10 mM (pH 7.0) as eluent A and Acetonitrile:Eluent A (95:5) as eluent B.

After purification, CoA-fluorophores were lyophilized and stored at -20°C in the dark. Before use, they were reconstituted in PBS and the concentration was determined through absorbance measurements.

#### *Production of Sfp*

Production and purification of 4'-phosphopantetheinyl transferase Sfp was performed according to the protocol in [5] with minor changes. Briefly, the bacterial suspension expressing the DNA plasmid coding for Sfp was centrifuged (6000 g, 4°C for 20 min) and the obtained pellet was resuspended in 10 mL of lysis buffer composed by 20 mM Tris-HCl at pH 8, 300 mM NaCl, 30 mM imidazole supplemented with protease inhibitor tablets (cOmplete™, EDTA-free Protease Inhibitor Cocktail, Sigma Aldrich), 0.05% of TritonX and 1 µg/ml of lysozyme. This suspension was sonicated on ice (six pulses of 30 s, separated by 60-s pauses), centrifuged (13000 g, 4°C for 30 min) and finally filtered (0.45-μm syringe filter). Purification was performed using a gravity-flow column HisPur™ Ni-NTA Resin (88221, Thermo Fisher Scientific) following the protocol for His-tagged proteins. The purified fraction was stored at -80°C in a solution of 20 mM Tris-Hcl at pH 7.5, 150 mM NaCl and 25% of glycerol.

#### *Cell culture, receptors expression and labelling*

SH-SY5Y (ECACC 94030304) and SK-N-BE(2) (ATCC® CRL-2271™) cell lines were grown at 37°C, 5% CO_2_, in DMEM/F-12 medium supplemented with 10% Fetal Bovine Serum, 1% Penicillin-Streptomycin, 1% L-Glutamine.

For live-cell experiments, cells were seeded in WillCo dishes or dishes with N-PK51 cover glasses (glued in the WillCo dishes after removing the original bottom) as bottom.

In the second case, we sterilized N-PK51 cover glasses by washing with ethanol and rinsing with Milli-Q water, followed by UV irradiation for two hours in a cell hood.

After 24 h from seeding, cells were transfected with S6-tagged human TrkA (SH-SY5Y) or p75^NTR^ (SK-N-BE(2)) constructs [6]. S6-p75^NTR^ was inserted in a lentiviral transfer plasmid as previously described [7], and this was used as is without reconstruction of lentiviral particles; S6-TrkA was obtained by cloning the human cDNA of S6-TrkA described previously [8] into a pcDNA3.1. Transfection was performed using Lipofectamine 2000 (Thermo Fisher Scientific) according to the manufacturer’s instructions. In the case of p75^NTR^, the expression was driven by an inducible promoter bearing a Tet-responsive element (TRE) [7]; therefore, 5 h after transfection, the expression was induced with 0.005 μg/ml of doxycycline added in the culture medium.

After 24-36 h, cells were labelled with a mix containing 2 μM Sfp Synthase, 10 mM MgCl_2_ and the CoA-conjugated form of the dye in the culture medium. We used 20nM of CoA-Atto 488 and 5 nM of CoA-Atto 565 for two-channel experiments; 5nM of CoA-Atto 565 or 5 nM of CoA-Abberior Star 635p for single-color experiments. The mix was added to cells for 15 min at 37°C, then they were washed five times with PBS and imaged in FluoroBrite DMEM supplemented with 0.5% BSA, 2mM of Trolox (Sigma-Aldrich) and 5μM of n-propyl gallate (Sigma-Aldrich).

#### *D 263® M glass autofluorescence*

Autofluorescence of WillCo dish cover glass made of Schott D 263® M was measured in a Renishaw micro-Raman system (Fig. 1C), with laser excitation at 473 nm, grating of 2400 line/mm, slit of 65 micron (corresponding to a spectral resolution of 1 cm^-1^) and focus within the glass.

#### *Backgrounds from different cover glass materials, cell media and immersion oils*

We compared the background observed using the following cover glass materials (Figure 1A, 2A): WillCo-dish® Glass Bottom dishes (WillCo Wells, cover glass made of Schott borosilicate glass D 263® M); MatTek glass bottom dishes (MatTek Life Sciences); FluoroDish™ glass bottom Cell Culture Dish (WPI World Precision Instrument); custom cover glasses made of Schott Phosphate Crown glass N-PK51 with the following specifications: thickness: 0.17mm +/-0.01mm, surface quality: 40/20, roughness: <1nmRMS, parallelism: 20 arcsec. Custom cover glasses were glued with a silicon glue below the WillCo dishes after removing the original bottom glass.

All the dishes were filled with PBS. The measurements were performed in TIRF with a penetration depth of 110 nm, using the Dual View system detailed above. All the laser powers (488 nm, 561 nm and 635 nm) were set to 1.4 mW after the objective without oil.

We also measured the background observed using the following immersion oils (Figure 3): Leica type F, Zeiss 518 F, Nikon type F, Cargille LDF and Olympus type F using a WillCo dish filled with PBS. We measured in TIRF mode (penetration depth: 110 nm) using simultaneous double excitation at 488 nm (3.5 mW at the objective) and 561 nm (3 mW), with corresponding simultaneous double detection.

To compare the background produced from cell media (Figure 5), WillCo dishes were filled with each one of the tested media: PBS; an imaging buffer suitable for living cells composed of 137 mM NaCl, 5.4 mM KCl, 2 mM CaCl_2_, 1 mM MgCl_2_, 10 mM HEPES, pH 7.5 [9]; RPMI 1640 Medium; DMEM medium; DMEM/F-12 medium; FluoroBrite™ DMEM medium. PBS, HEPES and cell media were purchased from Thermo Fisher Scientific; salts were purchased from Sigma Aldrich. To test the effects of common supplements as well (Figure S4), we added to the imaging buffer 1mM sodium pyruvate, 10% Fetal Bovine Serum (FBS), 0.5% bovine serum albumin (BSA), 4.5g/L D-Glucose, or 2mM L-glutamine. Moreover, we compared DMEM/F-12 without supplements, supplemented as a complete medium for cell growth (by adding 10% FBS and 2mM L-glutamine), supplemented as serum-starvation medium (by adding 0.5% BSA and 2mM L-glutamine). Measurements were performed in TIRF with single excitation at 488 nm (laser power at the objective 3.5 mW) and 561 nm (3mW), with a penetration depth of 110 nm.

In all background measurements, we acquired 100-frame time series with an integration time of 40 ms in different fields of view of each sample. We calculated the intensities averaged on the time series, subtracting the offset value.

#### *Measurements of chromatic aberration*

As chromatic aberration, we measured the focus shift in different channels using different immersion oils (Figure 3 A, B, C). We used TetraSpeck™ Microspheres (Invitrogen; ThermoFisher Scientific) with a diameter of 100 nm adhered on the bottom of the WillCo dish with its original D 263® M cover-glass thin-bottom or with the custom one made of N-PK51 Schott glass. We prepared samples at microspheres density low enough to distinguish all the single microspheres. The starting suspension containing ~1.8 × 10^11^ particles/mL was diluted 10 times in milliQ water; 5 μL of the diluted solution were deposited on the surface of the dish bottom and spread with the pipette tip. After waiting for the droplet to dry, the dish was filled with PBS for imaging. We acquired a z-stack in TIRF on different fields of view using simultaneous double excitation at 488 nm and 561 nm with corresponding simultaneous double detection. Measurements were performed at 37°C. Laser powers were adjusted to have similar SNR in the two channels.

For each bead simultaneously detected in the two channels, we extracted the dependence on the focus position of the average intensity within a fixed ROI containing the sphere image in each channel with ImageJ software (“z-profile”) and calculated the distance between the z-position of the intensity maximum in the two channels.

#### *Cell autofluorescence*

We prepared samples of SK-N-BE(2) cells at low confluence so that cell contours were distinguishable from areas without any cells around them. For each cell, we acquired a differential interference contrast (DIC) image and TIRF images with all combinations of excitation and detection reported in Figure 4. All laser powers (488, 561 and 635 nm) were set to 2mW at the objective. Integration time for TIRF images was 40 ms; penetration depth was varied from 100 to 190 nm (as determined by the Leica TIRF software considering the different excitation wavelengths), as reported in Figure 4.

Thanks to the DIC image, we identified the contours of the basal membrane for each cell. We measured the mean intensity inside these contours ($I_{in}$) and outside them in areas without any cells ($I_{out}$). We reported the relative increase of background caused by cell autofluorescence $r=\frac{I_{in}- I_{out}}{I_{out}}$.

#### *Fluorophore photostability in various solutions*

Trolox solutions were prepared by dissolving (±)-6-Hydroxy-2,5,7,8-tetramethylchromane-2-carboxylic acid (Sigma Aldrich) overnight at room temperature at a final concentration of 5mM. Solutions were prepared in PBS for experiments on fluorophores adhered to cover glass or in FluoroBrite DMEM for experiments in living cells. Trolox concentration was assessed by absorbance measurements using a UV/vis spectrophotometer (Jasco V-550), exploiting its known extinction coefficient of 2350 M^-1^ cm^-1^ in aqueous buffer at its absorption peak around 290 nm. Trolox quinone was generated by UV-irradiation (UV lamp Spectroline ENF 260C/FE, 6W, wavelength < 254 nm) and its concentration was determined from the absorbance at 255 nm with the following relationship, as previously described [10] (Fig. S7):

$$\left[ TQ \right]=\frac{\frac{A_{255}}{d}-\varepsilon_{255}\left( TX \right)\cdot{[TX]}_{0}}{\varepsilon_{255}\left( TQ \right)-\varepsilon_{255}(TX)}$$

where $A_{255}$ is the absorbance measured on a sample containing Trolox (TX) and Trolox quinone (TQ); ${[TX]}_{0}$ is the starting concentration of Trolox before oxidation; $\varepsilon_{255}\left( TX \right)=400 l \cdot{mol}^{-1}\cdot{cm}^{-1}$; $\varepsilon_{255}\left( TQ \right)=11600 l \cdot{mol}^{-1}\cdot{cm}^{-1}$, *d* id the optical path equal to 1 cm.

N-propyl gallate solutions were prepared by dissolving propyl gallate (Sigma-Aldrich) in PBS overnight at room temperature in the dark at a final concentration of 1mM.

Ascorbic acid solutions were prepared by dissolving L-Ascorbic acid (Sigma-Aldrich) in milli-Q water at a final concentration of 10mM; the pH was adjusted to 7.5 with NaOH.

All antibleaching solutions were used fresh within one day.

We measured bleaching on samples composed of a layer of fluorophores adhered to the bottom of Willco dishes (Fig. 6, 7, S6). We did a polylysine coating on the cover glass to promote fluorophores adhesion. The coating was prepared using Poly-L-lysine solution 0.1 % (w/v) in H_2_O (Sigma Aldrich). 150 μL of this solution were deposited to cover the cover glass of a 12 mm-aperture Willco dish. After incubation at room temperature for 10-15 minutes, the coating solution was removed and the surface was rinsed with PBS. Then a solution of fluorophores (typical concentration of about 1nM) in PBS was deposited on the coated cover glass for 10-30 minutes. After extensive washing to remove fluorophores remained in suspension or weakly adhering, the dish was filled with PBS or with antibleaching solutions and the sample was immediately observed with the TIRF microscope. We acquired time series (40 ms integration time, 56 ms frame time) to measure the temporal decay of emitted fluorescence. The duration of the time series (1000-8000 frames) was chosen in order to reach the plateau in the intensity profile in each condition. We used 3.5mW laser power for 488 nm excitation and 3mW for 561 nm excitation. For each sample, we acquired various fields of view.

The intensity decay was fitted with a bi-exponential function: $A_{1}\exp\left( -t/\tau_{1} \right)+A_{2}\exp\left( -t/\tau_{2} \right)+y_{0}$, in OriginLab (Origin Pro 2016), using a global fitting with a shared offset $y_{0}$for groups of measurements acquired on the same sample in the same condition in order to reduce correlation among fit parameters. We extract the decay time mediated with amplitudes: $\tau={(A}_{1}\tau_{1}+A_{2}\tau_{2})/{(A}_{1}+A_{2})$.

#### *Simulations*

We simulated spots diffusing with Brownian motion on a surface of 16 μm x 16 μm (density: 0.3 spot/ μm^2^) with a diffusion coefficient D of 0.55 μm^2^/s. Starting from randomly distributed spot positions, for each time step *dt* we calculated the displacement along x and y dimensions by using a gaussian distribution with mean 0 and standard deviation $\sqrt{2Ddt}$. The surface had reflective boundaries. For each simulation, the total time was 14 s, with a simulation time step of 56 ms, the pixel size was 0.16 μm and the standard deviation of each spot Gaussian point spread function was 1.2 pixels. A constant background and Gaussian noise were added to each pixel; an additional Shot noise was simulated by substituting each pixel value with a number generated using a uniform distribution (within an interval of 150) around a number calculated by multiplying by 150 a random number having a Poissonian distribution with average the original pixel value divided by 150. Spot intensity, background intensity, standard deviation of Gaussian noise were varied, and the value of 150 for the Shot noise was chosen empirically, in order to reproduce the experimental conditions (Table S4). These movies were created using MATLAB (The Mathworks, Inc.), in particular the functions randn, rand and poissrnd. We performed 5 different simulations for each condition. Each simulated movie was analyzed as described above for single particle tracking.

#### *Statistics*

Statistical analysis was performed with OriginPro 2016. We used the parametric tests Student’s t-test (two-tailed) or one-way ANOVA with Bonferroni or Tukey’s correction for multiple comparisons. When requirements for parametric tests were not met, we performed Kruskal–Wallis tests, followed by Dunn’s test for pairwise comparison with Bonferroni correction. Significance was set at α=0.05. Details for each applied test are provided in the figure captions.

### **Supplementary Figures**

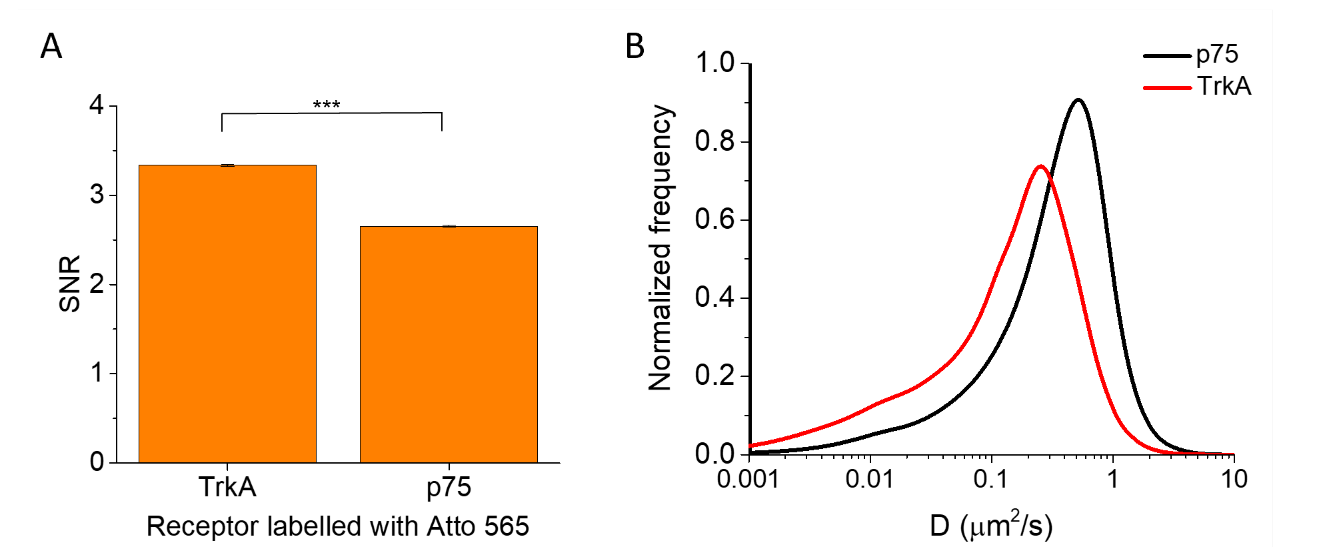

**Figure S1. Diffusivity impact on SNR.** A) SNR measured on TrkA and p75^NTR^ receptors labeled on the membrane of living cells with Atto 565. Data were extracted from acquisitions in a two-color TIRF setup including 488 nm excitation. Results are mean ± SEM, obtained from 5 movies on different cells from 2 independent repetitions, with 1500-3500 spots analyzed in each movie. *** P < 0.001, Student's t-test. B) Distribution of diffusion coefficient (D) for p75^NTR^ (black curve, 403 trajectories from 4 cells from 2 independent repetitions) and TrkA receptors (red curve, 603 trajectories from 7 cells from 2 independent repetitions), estimated with single-particle tracking.

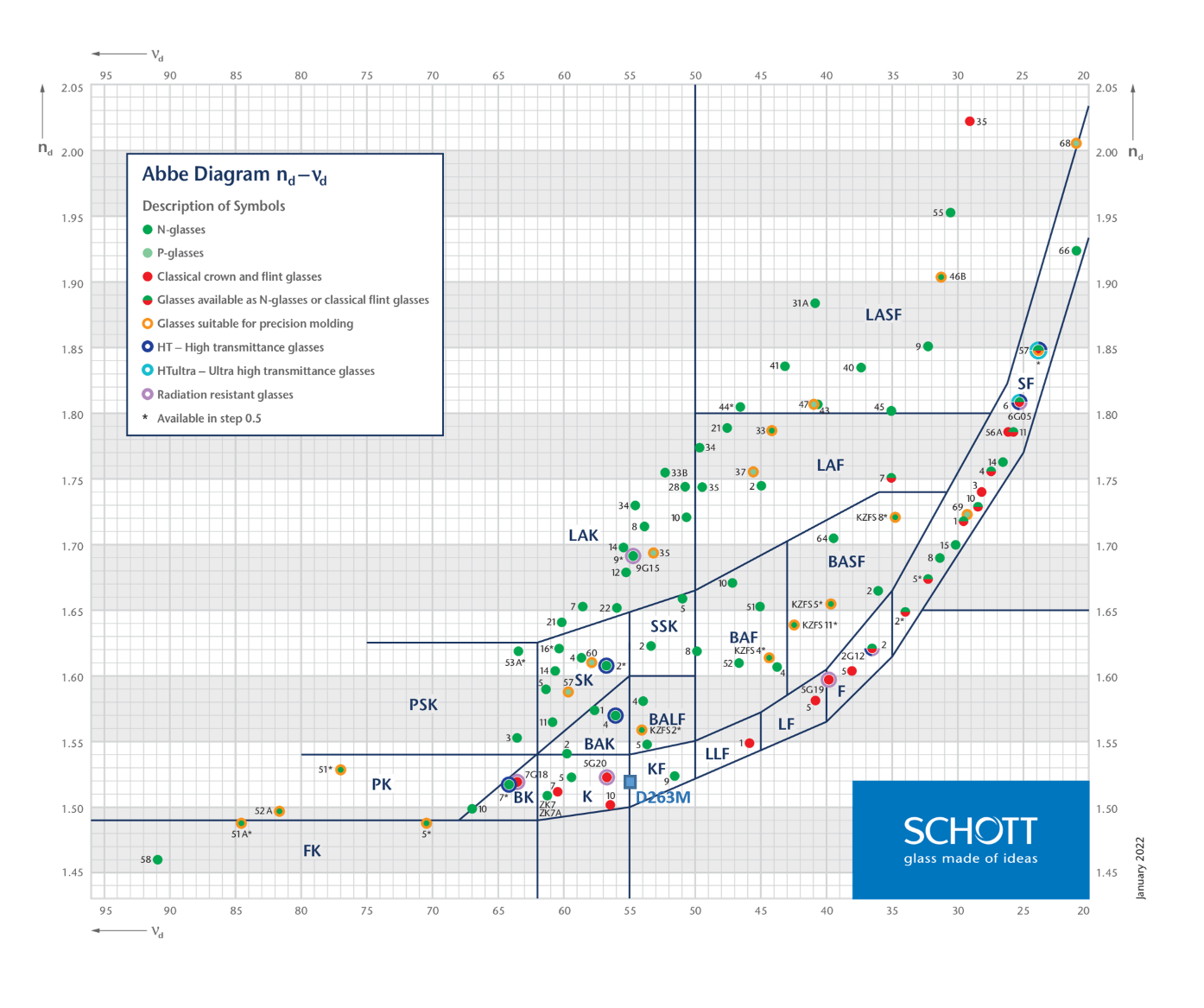

**Figure S2. Abbe diagram for Schott glasses**. The diagram plots the refractive index at 587.56 nm (n_d_) versus the Abbe number (ν_d_) for glasses produced by Schott company. Glass families (separated by lines on the graph) are identified by letter codes depending on the main composition; inside each family, each glass (circles on the graph) is identified by an additional alphanumerical code. Figure reproduced with permission from Schott website [11] (© SCHOTT AG, Optical Glass Abbe Diagram) with addition of the blue square corresponding to reported properties for D263M cover glasses. N-glasses: environmentally friendly alternative to conventional lead and arsenic-containing glass types; P-glasses: low transition temperature glasses for precision molding process.

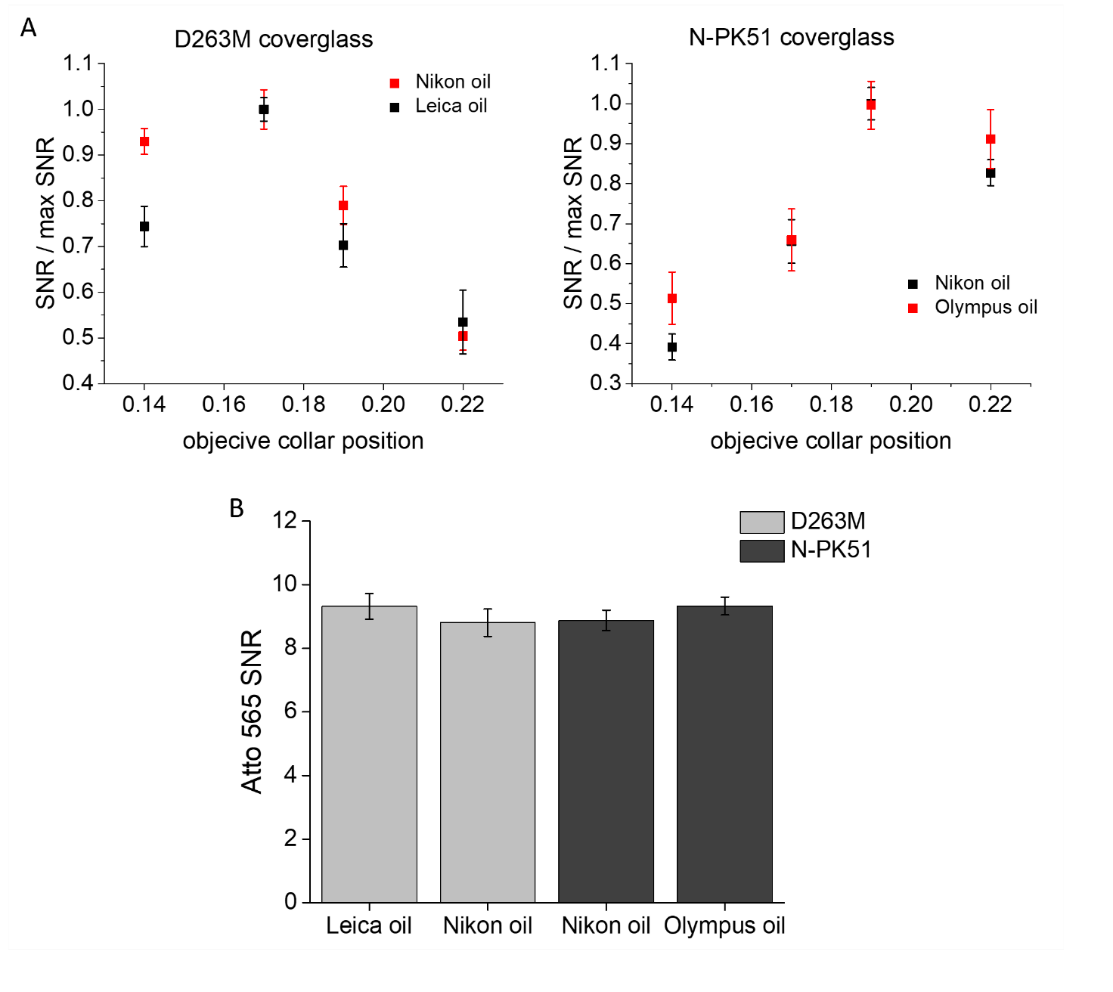

**Figure S3. Effects of immersion oil on SNR.** A) SNR measured in TIRF for single molecules of Atto 565 as a function of objective collar position using two different immersion oils. The SNR is normalized to the maximum observed value. Left: D263M cover glass; right: N-PK51 cover glass. Results are mean ± SEM, obtained, for each condition, from 5 movies in different fields of view from 2 independent repetitions, with 2000-20000 spots analyzed in each movie. B) SNR (mean ± SEM) of Atto 565 single dyes adherent to the cover glass measured with objective collar positions corresponding to the maximum SNR. Differences are not significant according to an ANOVA test.

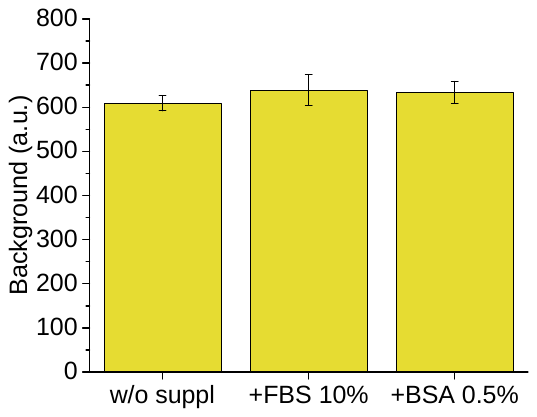

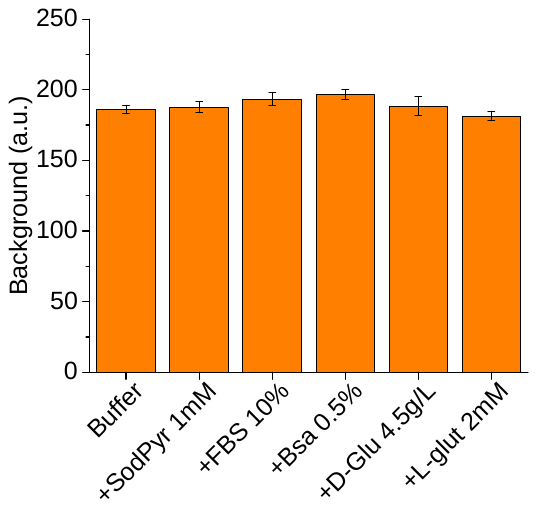

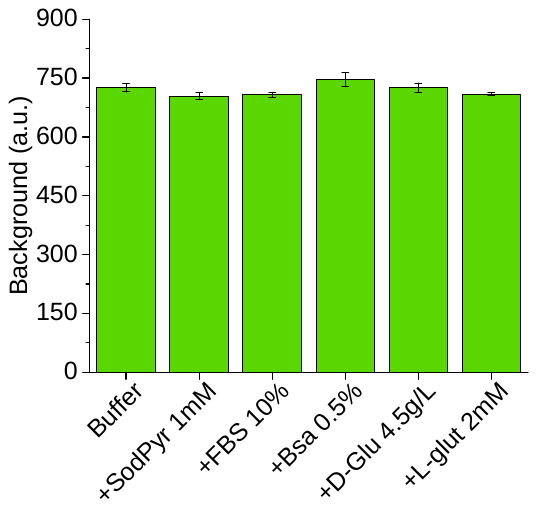

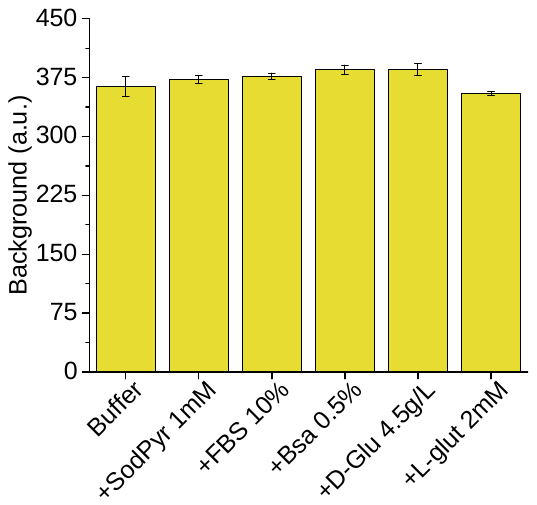

**Exc: 488 nm, det: green**

**Exc: 488 nm, det: orange**

**Exc: 561 nm, det: orange**

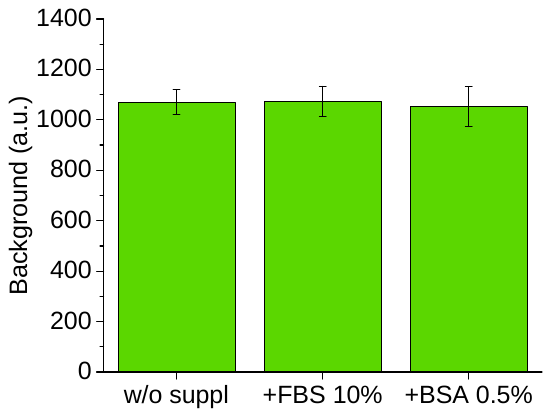

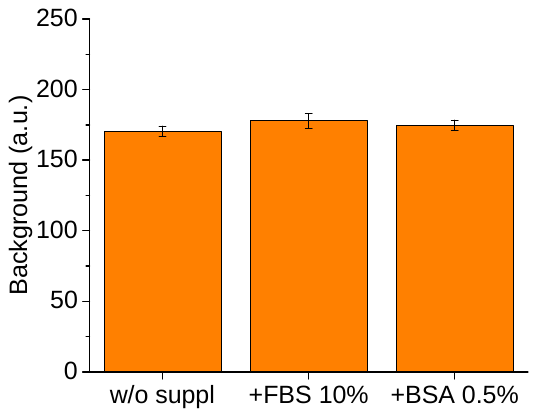

DMEM/F-12

DMEM/F-12

DMEM/F-12

A

B

w/o suppl

+ FBS 10%

+ L-Glu 2mM

+ BSA 0.5%

+ L-Glu 2mM

w/o suppl

+ FBS 10%

+ L-Glu 2mM

+ BSA 0.5%

+ L-Glu 2mM

w/o suppl

+ FBS 10%

+ L-Glu 2mM

+ BSA 0.5%

+ L-Glu 2mM

**Figure S4. Autofluorescence for different media supplements**. A) Comparison of background detected in TIRF when supplements are added in a buffer composed of HEPES and salts (see Material and Methods for its detailed composition). B) Comparison of background detected in TIRF for DMEM/F-12 medium without supplements, supplemented as complete growth medium (FBS 10%, L-Glu 2mM) and supplemented as serum-starvation medium (BSA 0.5%, L-Glu 2mM). Excitations and detections are as in panel A. Results are mean ± SEM, obtained from 6-12 different fields of view from two independent repetitions. Differences are not significant in ANOVA testing.

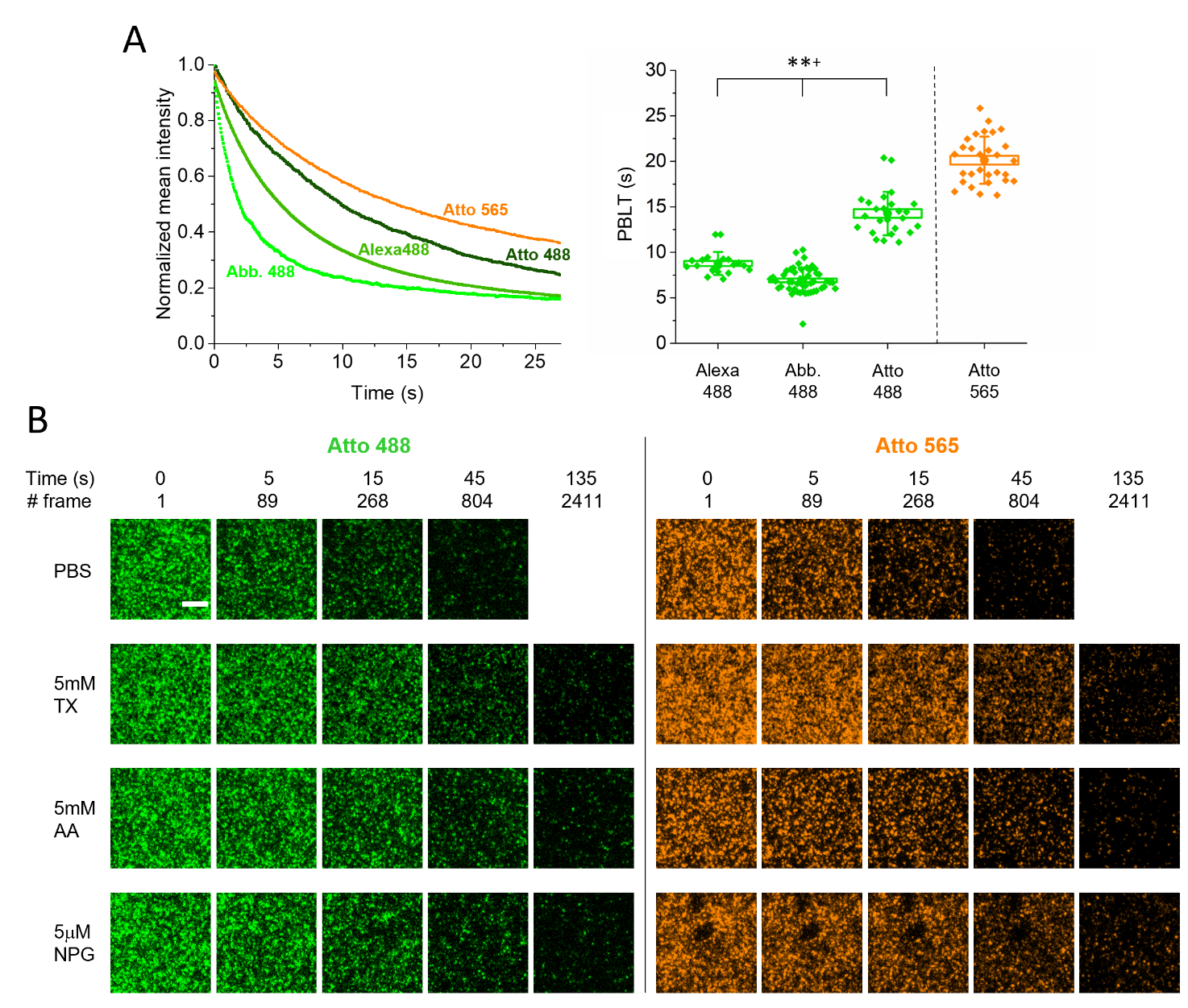

**Figure S5. Comparison of dyes photobleaching**. A) Left: examples of temporal decay of intensity observed in TIRF for the different indicated dyes. Abb. 488: Abberior STAR 488. Right: extracted photobleaching lifetimes (PBLTs) of Alexa 488, Abb.488 and Atto 488 are compared. **^+^ P < 0.002, Kruskal Wallis, Dunn's means comparisons with Bonferroni correction. Atto 565 lifetime is also reported. Box: SEM, whiskers: standard deviation, spots: individual measurements from two independent repetitions. B) Atto 488 and Atto 565 imaged in TIRF using PBS medium or PBS supplemented with Trolox (Tx), Ascorbic Acid (AA) or n-propyl gallate (NPG). Representative images from the acquired time series (frame time: 56 ms). Scale bar: 5μM.

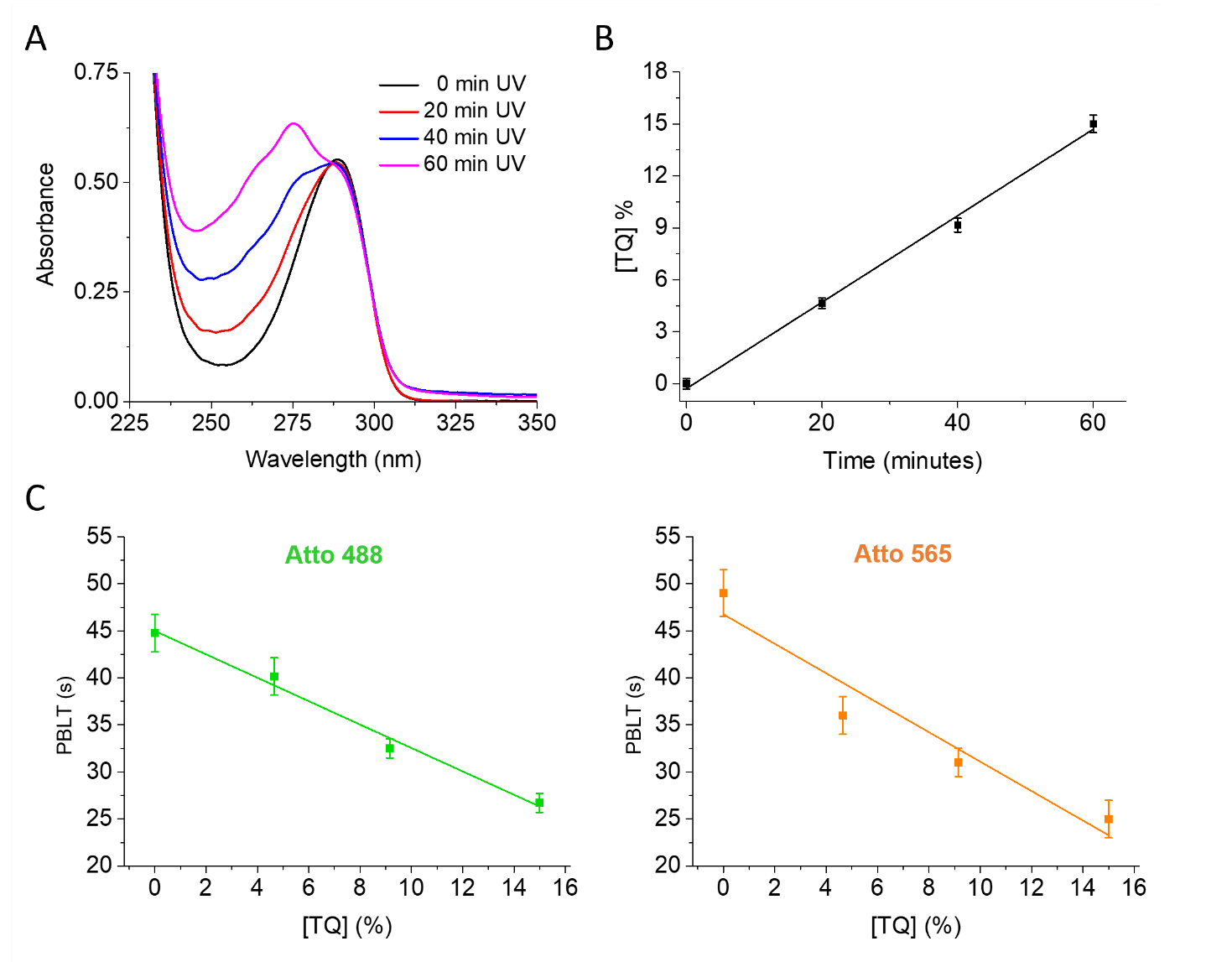

**Figure S6. Trolox oxidation.** A) Oxidation of Trolox was induced by UV light using different illumination times and was monitored by absorbance measurement. It is possible to observe: the characteristic peak of Trolox at ~290 nm, the formation of the new peak at ~270 nm, corresponding to the Trolox-quinone, the isosbestic point at ~285 nm. B) Percentage of Trolox-quinone concentration [TQ], calculated from the absorbance measures as described in Material and Methods, plotted against UV illumination time. Data from two independent repetitions. C) Effect of Trolox oxidation on Atto 488 (left, green) and Atto 565 (right, orange) photobleaching lifetimes (PBLTs) observed in TIRF. Measurements were performed using 2mM of Trolox/TQ mixtures with different percentages of TQ. Data are obtained from 6-12 fields of view from two independent repetitions. All experimental points are mean ± SEM and continuous lines are linear fitting curves.

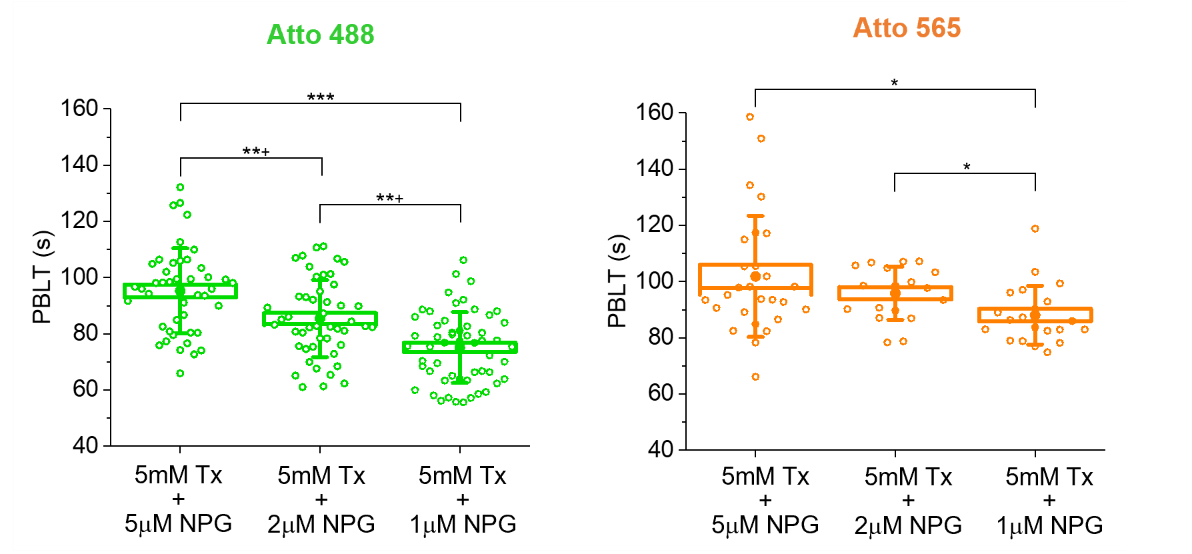

**Figure S7. Photobleaching lifetime with mixtures of Trolox and n-propyl gallate.** Photobleaching lifetimes (PBLTs) observed by mixing three concentrations of n-propyl gallate (NPG) with 5mM of Trolox (TX). *** P < 0.001, **^+^ P < 0.002, 1-way ANOVA, Bonferroni multiple comparisons. * P < 0.05, Kruskal Wallis, Dunn's means comparisons with Bonferroni correction. Results are reported for Atto 488 (left, green) and Atto 565 (right, orange). Box: SEM; whisker: SD; dots: individual fields of view, from 2-5 independent repetitions.

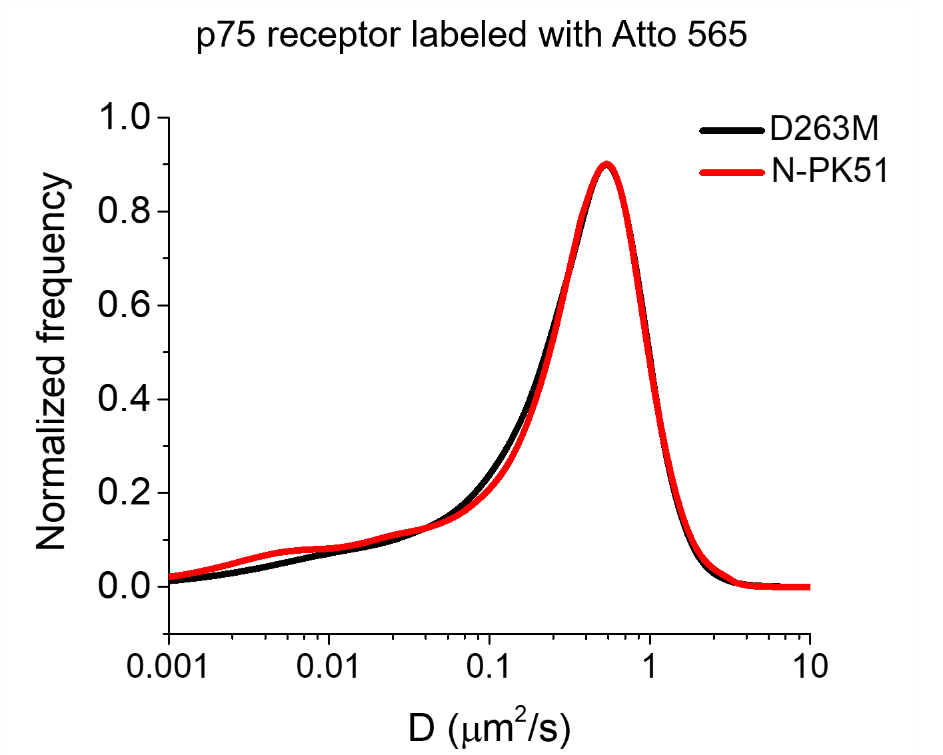

**Figure S8. Single particle tracking of p75 receptors on D263M and N-PK51 cover glasses**. The distribution of diffusion coefficients (D) for p75 receptors was estimated through single particle tracking analysis on single channel TIRF movies. Receptors were labelled with Atto 565 on the membrane of SK-N-BE(2) cells seeded on D263M cover glass (black, 4086 tracks) and N-PK51 cover glass (red, 2127 tracks). Trajectories were obtained from 6 cells from two independent repetitions in each case.

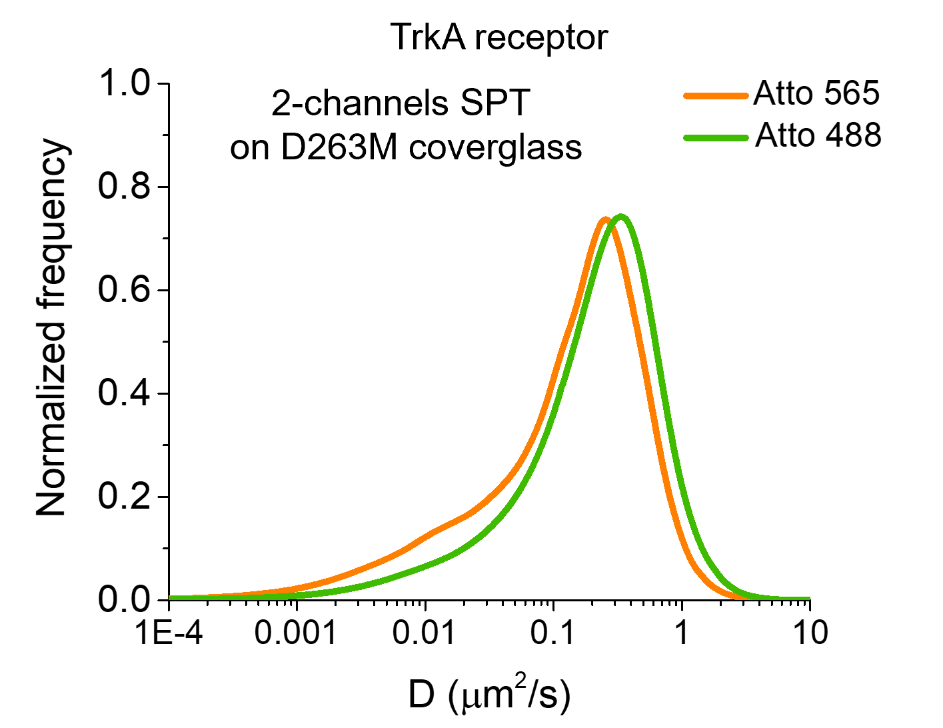

**Figure S9. Two-color single particle tracking of TrkA receptors on D263M cover glass**. Distributions of diffusion coefficients (D) for TrkA receptors expressed in SH-SY5Y cells labelled simultaneously with Atto 565 (orange curves) and Atto 488 (green curves). D was estimated through single particle tracking analysis on simultaneous two-color TIRF acquisitions. Atto 565: 2973 tracks. Atto 488: 2594 tracks. Trajectories were obtained from 8 cells from two independent repetitions.

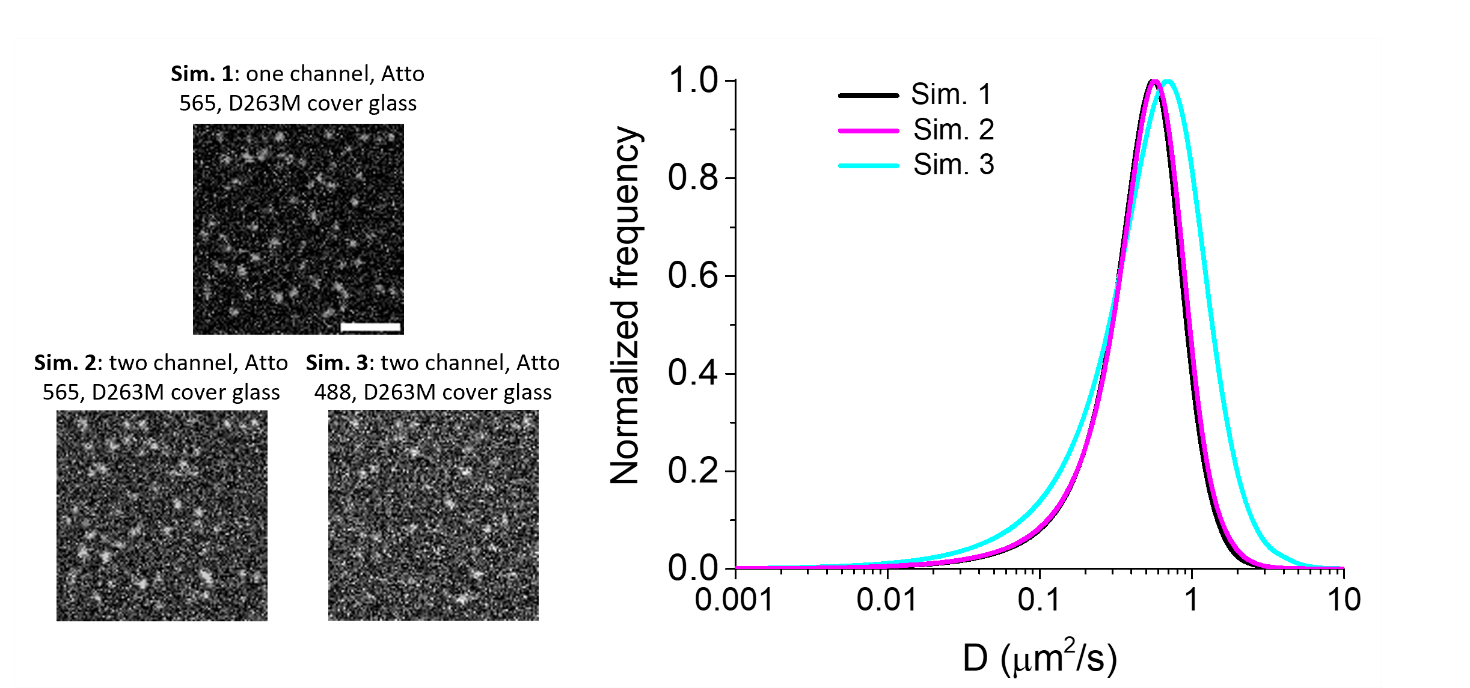

**Figure S10. Single particle tracking on simulations reproducing conditions observed on D263M cover glass.** Left: examples of images from the simulated movies (showed with the same grayscale). Experiments with similar signal intensities, background and noise are indicated for each simulation. Scale bar: 5μm. Right: distributions of the diffusion coefficient (D) estimated with single particle tracking analysis on the simulated movies. Sim. 1: 2458 tracks, Sim. 2: 2178 tracks, Sim. 3: 2280 tracks; obtained in each case from 5 different simulations. Histogram maxima are normalized to 1.

### **Supplementary Tables**

| **Dye** | **Extinction coefficient (M^-1^ cm^-1^)** | **Quantum yield (%)** | **Brightness (M^-1^ cm^-1^)** |
| --- | --- | --- | --- |
| Abberior STAR 635p | 120000 | 90 | 108000 |
| Atto 565 | 120000 | 90 | 108000 |
| Alexa 488 | 72000 | 92 | 66240 |
| Atto 488 | 90000 | 80 | 72000 |
| Abberior STAR 488 | 65000 | 89 | 57850 |

**Table S1**. Properties of the dyes used in the study, as reported by manufacturers.

| **Cover glass** | **Refractive index n_e_** | **Refractive index n_D_** | **Thickness (mm)** | **Abbe number** |
| --- | --- | --- | --- | --- |
| D263M | 1.5255 | 1.5230 | 0.170 ± 0.005 | 55 |
| N-PK51 | 1.5302 | 1.5286 | 0.170 ± 0.01 | 77 |

**Table S2**. Properties of the cover glasses used in the study. First row: cover glasses made of standard D263M material (present as glass-bottom in WillCo-Dishes); second row: custom cover glasses made of N-PK51 material. Refractive indexes are reported at 587.56 nm (n_D_) and at 546.07 nm (n_e_).

| **Immersion oil** | **Refractive index n_e_ (23 °C)** | **Refractive index n_D_ (23 °C)** | **Viscosity** | **Abbe number ν_e_ (23 °C)** | **Density** |
| --- | --- | --- | --- | --- | --- |
| Leica type F | 1.518 | 1.515 | 500 mm²/s at 20°C  435 mm^2^/s at 23 °C | 45.8 | 1,093 g/cm^3^ at 20°C |
| Nikon type F | 1.518 | 1.515 | 410 mm²/s at 23 °C | 40.8 | 0.8-1.0 g/cm^3^ at 20°C |
| Olympus type F | 1.518 | 1.515 | 450 mm^2^/s at 23 °C | 40.8 | 0.9169 g/cm^3^ at 15 °C |
| Zeiss 518 F | 1.518 | 1.515 | 445 mm²/s at 20 °C | 45 | 1,093 g/cm^3^ at 20 °C |
| Cargille LDF | 1.518 | 1.515 | 500 mm^2^/s at 23 °C | 41.3 | 0.984 g/cm^3^ at 23 °C |

**Table S3**. Properties of the immersion oils used in the study, as reported by manufacturers. Refractive indexes are reported at 587.56 nm (n_D_) and at 546.07 nm (n_e_).

| **Parameter** | **Sim. 1** | **Sim. 2** | **Sim. 3** |
| --- | --- | --- | --- |
| Spot intensity (±30%) | 1350 | 1350 | 900 |
| Background intensity | 400 | 650 | 850 |
| Gaussian noise standard deviation | 200 | 300 | 350 |

**Table S4.** Simulation parameters varied to reproduce different experimental conditions (see also Fig. S10). Sim. 1 simulates one-color TIRF acquisition of p75^NTR^ receptors labeled with Atto 565 on D263M cover glass; Sim.2 and Sim. 3 simulates two-color TIRF acquisition of p75^NTR^ receptor labeled with Atto 565 (Sim. 2) and Atto 488 (Sim. 3) on D263M cover glass. To the spot intensity, we assigned the reported mean value with a normally distributed uncertainty of 30%.

### **Supplementary Videos**

**Supplementary Video 1**. Two-color single-molecule imaging of S6-p75 receptors moving on the membrane of living cells. Receptors were labeled simultaneously with Atto 488 (left channel) and Atto 565 (right channel). Cells were cultured on D263M cover glasses (glass-bottom of WillCo dishes). Scale bar: 5 μm.

**Supplementary Video 2**. Single-particle tracking analysis performed on the Atto 488 channel of Supplementary Video 1. A mask was applied to select the cell area. In green: trajectories reconstructed with u-track software. Scale bar: 5 μm.

**Supplementary Video 3**. Single-particle tracking analysis performed on the Atto 565 channel of Supplementary Video 1. A mask was applied to select the cell area. In red: trajectories reconstructed with u-track software. Scale bar: 5 μm.

**Supplementary Video 4**. Two-color single-molecule imaging of s6-p75 receptors moving on the membrane of living cells. Receptors were labeled simultaneously with Atto 488 (left channel) and Atto 565 (right channel). Cells were cultured on custom cover glasses made of N-PK51 material. Scale bar: 5 μm. Same grayscale as in Supplementary Video 1.

**Supplementary Video 5**. Single-particle tracking analysis performed on the Atto 488 channel of Supplementary Video 4. A mask was applied to select the cell area. In green: trajectories reconstructed with u-track software. Scale bar: 5 μm.

**Supplementary Video 6**. Single-particle tracking analysis performed on the Atto 565 channel of Supplementary Video 4. A mask was applied to select the cell area. In red: trajectories reconstructed with u-track software. Scale bar: 5 μm.
